# Bronze Age Northern Eurasian Genetics in the Context of Development of Metallurgy and Siberian Ancestry

**DOI:** 10.1101/2023.10.01.560195

**Authors:** Ainash Childebayeva, Fabian Fricke, Adam Benjamin Rohrlach, Lei Huang, Stephan Schiffels, Outi Vesakoski, Lena Semerau, Franziska Aron, Vyacheslav Moiseyev, Valery Khartanovich, Igor Kovtun, Johannes Krause, Sergey Kuzminykh, Wolfgang Haak

**Affiliations:** Department of Archaeogenetics, Max Planck Institute for Evolutionary Anthropology, D-04103 Leipzig, Germany; Department of Anthropology, University of Kansas, Lawrence, KS 66044, USA; German Archaeological Institute, Eurasia Department, Berlin 14195, Germany; School of Computer and Mathematical Sciences, University of Adelaide, Adelaide SA 5005, Australia; Department of Finnish and Finno-Ugric Languages, University of Turku, Turku 20014, Finland; Department of Archaeogenetics, Max Planck Institute for the Science of Human History, Jena 07745, Germany; Peter the Great Museum of Anthropology and Ethnography (Kunstkamera), Russian Academy of Sciences, University Embankment, 3, Saint Petersburg 199034, Russia; T.F. Gorbachev Kuzbass State Technical University, Department of History, Philosophy and Social Sciences, Kemerovo 650000, Russia; Russian Academy of Sciences, Institute of Archaeology, Laboratory of Natural Scientific Methods, Moscow 117292, Russia

**Keywords:** ancient DNA, Bronze Age Eurasia, Siberian Ancestry, population genetics

## Abstract

The Eurasian Bronze Age (BA) has been described as a period of substantial human migrations, the emergence of pastoralism, horse domestication, and development of metallurgy. This study focuses on individuals associated with BA metallurgical production, specifically the Seima-Turbino (ST) phenomenon (∼2,200-1,900 BCE) associated with elaborate metal objects found across Northern Eurasia. The genetic profiles of nine ST-associated individuals vary widely ranging between ancestries maximized in individuals from the Eastern Siberian Late Neolithic/BA, and those of the Western Steppe Middle Late BA. The genetic heterogeneity observed is consistent with the current understanding of the ST metallurgical network as a transcultural phenomenon. The new data also shed light on the temporal and spatial range of an ancient Siberian genetic ancestry component, which is shared across many Uralic-speaking populations, and which we explore further via demographic modeling using additional genome-wide (2 individuals) and whole genome data (5 individuals, including a ∼30x genome) from northwestern Russia.

## Introduction

Bronze Age Eurasia (∼3000-1000 BCE) is characterized by the development of metallurgy, one of the most important cultural innovations in human history. The Early Bronze Age in Eurasia (∼3000 BCE) is associated with the emergence of the Circumpontic Metallurgical Province, and eastward expansion of metallurgical production and exchange across the Eurasian steppe^1–3^. In the Late Bronze Age (∼2200–1000 BCE), a westward movement of materials was also detected, specifically in connection with the so-called Seima-Turbino (henceforth ST) phenomenon^1,2^ as seen by the presence of specific metal artifacts throughout the forest and forest-steppe regions of Northern Eurasia^4^.

The ST is represented by several sites throughout Eurasia dating to ∼2,200-1,900 BCE and constitutes a “metallurgical network” represented by many shared traits, such as the use of tin-copper, comparable artifact types, and shared metallurgical technologies that may have involved a movement of craft workers and/or groups^4,5^. The ST has been described as a “transcultural” phenomenon, i.e., a network of metallurgical production with shared traits on top of the underlying basis of consistent pottery types in the different areas associated with various archaeological cultures throughout northern Eurasia.

The name Seima-Turbino derives from the two eponymous burial grounds Seima and Turbino excavated in the beginning of the 20^th^ century CE^4^. The ST phenomenon combines elements of several cultures and does not represent a single culture in itself, especially since there are no distinct settlements or pottery styles associated with it. Instead, certain metal objects, which were found throughout Eurasia, from China and Central Asia in the East to Finland and Moldova in the West, spanning across approximately three million square kilometers, represent key ST-associated criteria (see Supplementary Note 1, Figure 1, and Supplementary Figure 1). In the entire spatial distribution of the ST, there is a certain degree of regional variation (see Supplementary Note 1). Briefly, the artifacts in the east contain higher amounts of tin (Sn) (Supplementary Figures 2 and 3) and more casting molds have been identified in the east compared to the west of the Urals, although a greater number and variation of ST objects have been found west of the Urals (Figure 1c). The metallic inventory of ST-complexes can be divided into two major groups: (1) objects that can be attributed to Eurasian archaeological cultures, such as Alakul, Abashevo, Sintashta, Petrovka and Srubnaya; (2) objects that are only known from ST-sites (socket axes, lamellar dagger blades, fully hilted daggers and knives and the so-called forked lanceheads) (Figure 1c). Singular finds of objects and weaponry of the ST type have been reported from contexts otherwise attributed to Bronze Age Glazkovo, Okunevo, Elunino, Odino, Krotovo, Koptyaki, Sintashta, Petrovka, Abashevo, Srubnaya (Pokrovka), and Post-Fatyanovo cultures^4,6^. However, these finds are rare and most ST materials are found at ST-associated sites.

**Figure 1.**
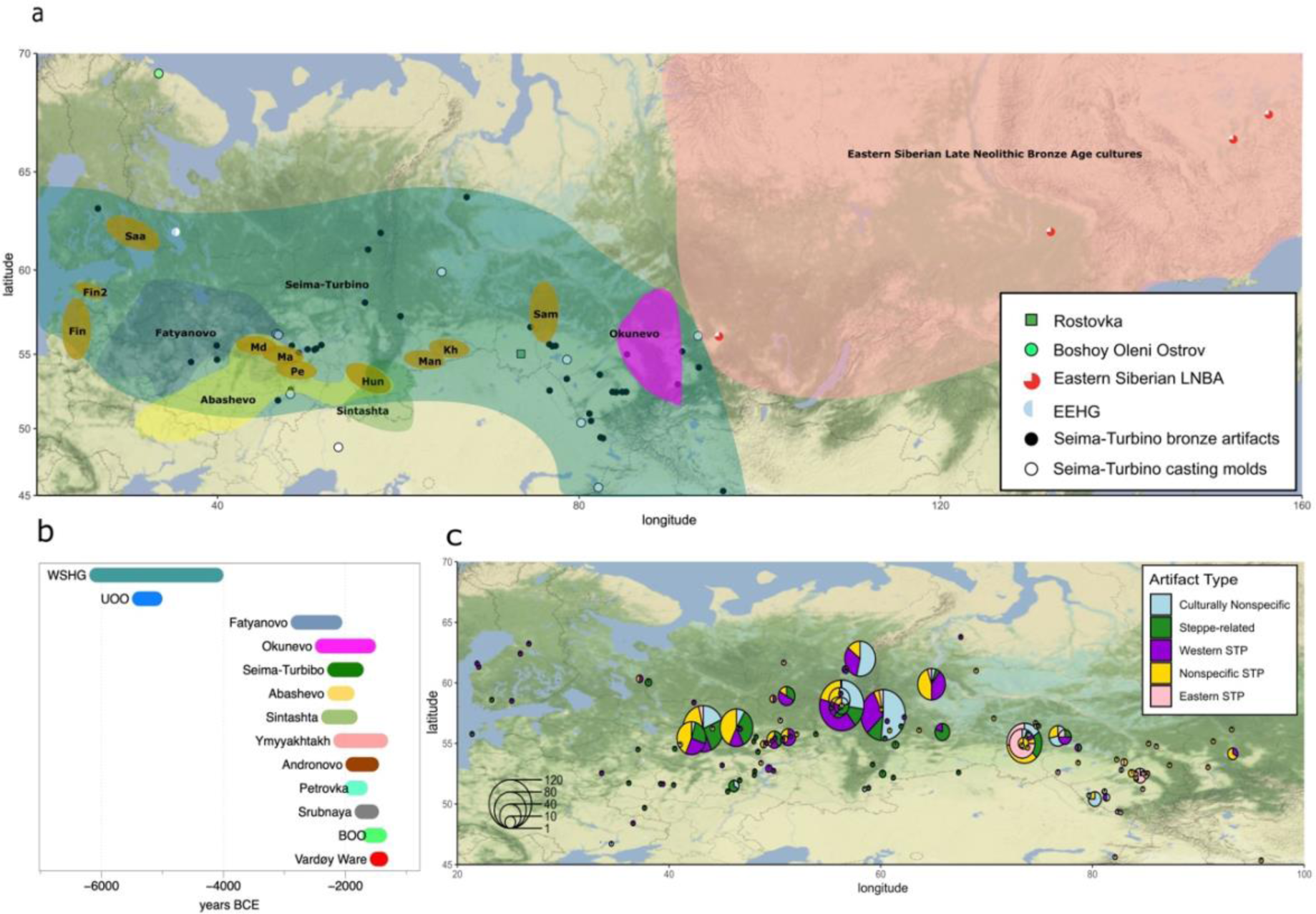
(a) Geographic map with ROT and BOO indicated, also showing hypothetical origin locations for ancestral stages of Uralic subfamilies (Saa=Saami, Fin/Fin2=Finnic, Man=Mansi, Kh=Khanty, Sam=Samoyedic, Hun=Hungarian, Md=Mordvin, Ma=Mari, Pe=Permic), and a distribution of contemporaneous archaeological cultures (adapted from Grünthal et al. 2022), (b) Chronology of Seima-Turbino (ST is including ROT) and BOO individuals together with relevant Bronze Age groups of Northern Eurasia. The timeline is based on a combination of absolute (^14^C) and relative dates, (c) Cultural/regional attribution of the metallic inventory of the sites of the ST phenomenon. Pie charts indicate the breakdown of artifacts at specific sites by cultural/regional attribution.

The people buried with ST-objects have been described as metallurgists who developed elaborate and distinct bronze objects, and possibly used river systems for transportation^4^. Even though the horse plays a central role in the ST iconography, it remains unclear whether people associated with the phenomenon were using horses for riding, traction or transport. It has been hypothesized that the number of people associated with the ST phenomenon was small, since there are very few sites with human remains linked to the phenomenon, and ST metal artifacts are comparably few but geographically widespread. ST burials are very distinct from those of the other North Eurasian cultures: individuals were buried mostly without pottery and not in kurgans, both inhumations and cremations were common, and the grave goods included bronze, stone, and bone weaponry, as well as bone armor. In cases where pottery is present at ST-sites, it can be attributed to other local cultures, for example Koptyaki at the site Shaitanskoe Ozero II^7^. The early history of the ST phenomenon is not well understood, however, based on the presence of tin and copper in metal alloys of ST objects, the Altai and Tian-Shan mountains have been proposed^4,8^.

Here, we present ancient human DNA data from a well-known, ST-associated site Rostovka (ROT), which is one of the very few ST sites with preserved human remains. The majority of the graves found at Rostovka contain bronze ST objects, bronze weapons and tools, casting molds, jewelry, bone knife handles, and armor plates^9^. In fact, the majority of Rostovka burials contain weapons, found in 60% of the female and 80% of the male burials^10^. The radiocarbon dates for Rostovka, excluding extreme values, range between ca. 2200-2000 cal. BCE^3,9,11^. This chronological horizon is simultaneous with the Okunevo culture, while its lower bound overlaps with the early Abashevo and Sintashta cultures, among others (Fig. 1b).

An identifying component in the genetic landscape of Northern Eurasia is a shared Siberian ancestry component, which is present in the genetic profiles of Finnish, Estonian, Saami-speaking, and indigenous Siberian populations today^12^. A previous ancient DNA study focusing on the Eastern Baltic found a genetic contribution from Siberia in the Iron Age but not in the Bronze Age, which was linked to the time of the arrival of Uralic languages to the region^13^. Moreover, the Y-haplogroup N1a1a1a1a (previously known as N3a), primarily found in present-day northern Eurasian and Uralic speaking groups, first appears in Europe in Early Metal Age individuals from the Bolshoy Oleni Ostrov site (BOO)^14^, in northwestern Russia, together with evidence of high levels of genome-wide Siberian ancestry^15^.

We present new genome-wide data from two additional BOO individuals and shotgun data for five published individuals (including one high coverage genome of ∼31x). Direct or indirect contacts between BOO and southern and western Scandinavia have been proposed based on genetic data and the archaeological record^14–16^, but BOO has not been associated with any known early Metal Age cultures.

Here, we report the results of joint population genetic analyses of both sites in comparison with published ancient data from chronologically, geographically, and archaeologically relevant cultures of the forest-tundra (taiga and tundra) and forest-steppe zones of Eurasia. We also investigate the demographic history of Northern Eurasia, especially in context of the Siberian genetic component. Together, we aim to provide an updated view on the genetic history and connections of populations of the forest-steppe and western Siberia, with an emphasis on the ST phenomenon in the context of metallurgical production.

## Results

In this study, we report genome-wide SNP data for nine individuals from the ST site Rostovka, as well as additional new data for two BOO individuals (plus shotgun genome data for five already published individuals) (Fig. 1a). We performed 1240k SNP^17,18^ and mitochondrial genome captures on the nine individuals from ROT, and the two new BOO individuals, as well as Y-chromosomal capture^19^ on just the males. Lastly, we generated shotgun sequence data for five published BOO individuals, including one 31.8x covered individual (Fig. 1a, Table 1, Supplementary Table 1). Of the newly analyzed individuals, eight ROT individuals were genetically male and one was female, while both new BOO individuals were female (Table 1). Biological relatedness analysis of the newly reported individuals was performed using READ^20^ and lcMLkin^21^. Based on these analyses, we identified a pair of second-degree relatives (ROT011 and ROT015), both of whom are males carrying the Y-haplogroup C2a, and could either represent a grandson/grandparent, a nephew/uncle pair or paternal half-siblings, consistent with overlapping radiocarbon dates for both individuals (Table 1). A second-degree related pair was also found among the BOO individuals (BOO004-BOO005).

**Table 1.**
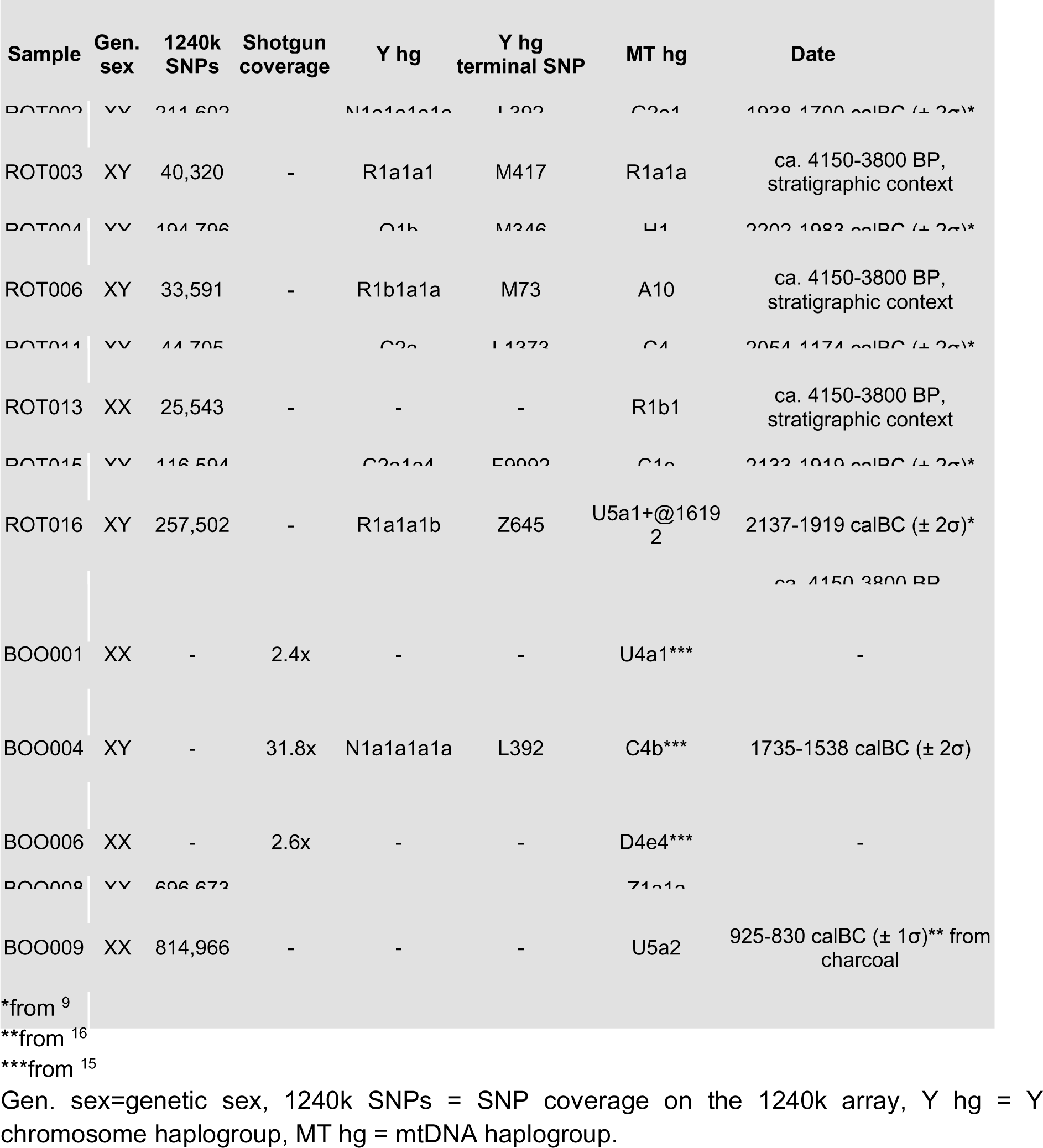
General overview of the ROT and BOO individuals included in the study.

We also generated a radiocarbon date for individual BOO004, whose genome was shotgun sequenced to 31.8x coverage (Table 1). The radiocarbon date (MAMS-57646) for this individual was determined to be 3351±25 BP, or 1735-1538 calBC (± 2σ) after calibration with OxCal 4.4^22^. However, we did not correct for a potential freshwater reservoir effect, which may place BOO004 at a younger date.

### General population genetic results

We used smartPCA^23^ to perform a principal component analysis (PCA) of modern-day reference populations from Eurasia and the Americas, onto which the ROT and the BOO individuals were projected (Fig. 2a and b). When assessing the genetic structure of Eurasian populations, plotting PC1 vs. PC2 (Fig. 2a) allows us to separate west and east Eurasian populations from the Native American groups, while plotting PC1 vs PC3 (Fig. 2b) distinguishes the major Eurasian ecological zones^24,25^. Looking at the newly generated data, ST individuals spread widely on the Eurasian PCA (PC1 vs PC3), mainly throughout the so-called ‘forest-tundra’ genetic cline (Fig. 2b) mirroring the distribution of the modern Uralic speakers (Supplementary Figure 4). Based on an unsupervised ADMIXTURE analysis^26^ of a reference set of published ancient data with K=7 clusters (Fig. 2c, Supplementary Figure 5), the ROT individuals generally carry diverse ancestry components ranging between a genetic profile represented by the Western Steppe Middle Late Bronze Age cluster (Western_Steppe_MLBA, we use Sintashta_MLBA when modeling this ancestry going forward)^27^ - a combination of orange and dark and light blue colors - and that of the Late Neolithic/Bronze Age East Siberians (Eastern_Siberia_LNBA)^28^ - red color (Fig. 2c). In comparison, the BOO individuals form a tighter and more homogeneous cluster that can be seen with both the PCA and the ADMIXTURE analyses, in line with what has been previously reported^15^.

**Figure 2.**
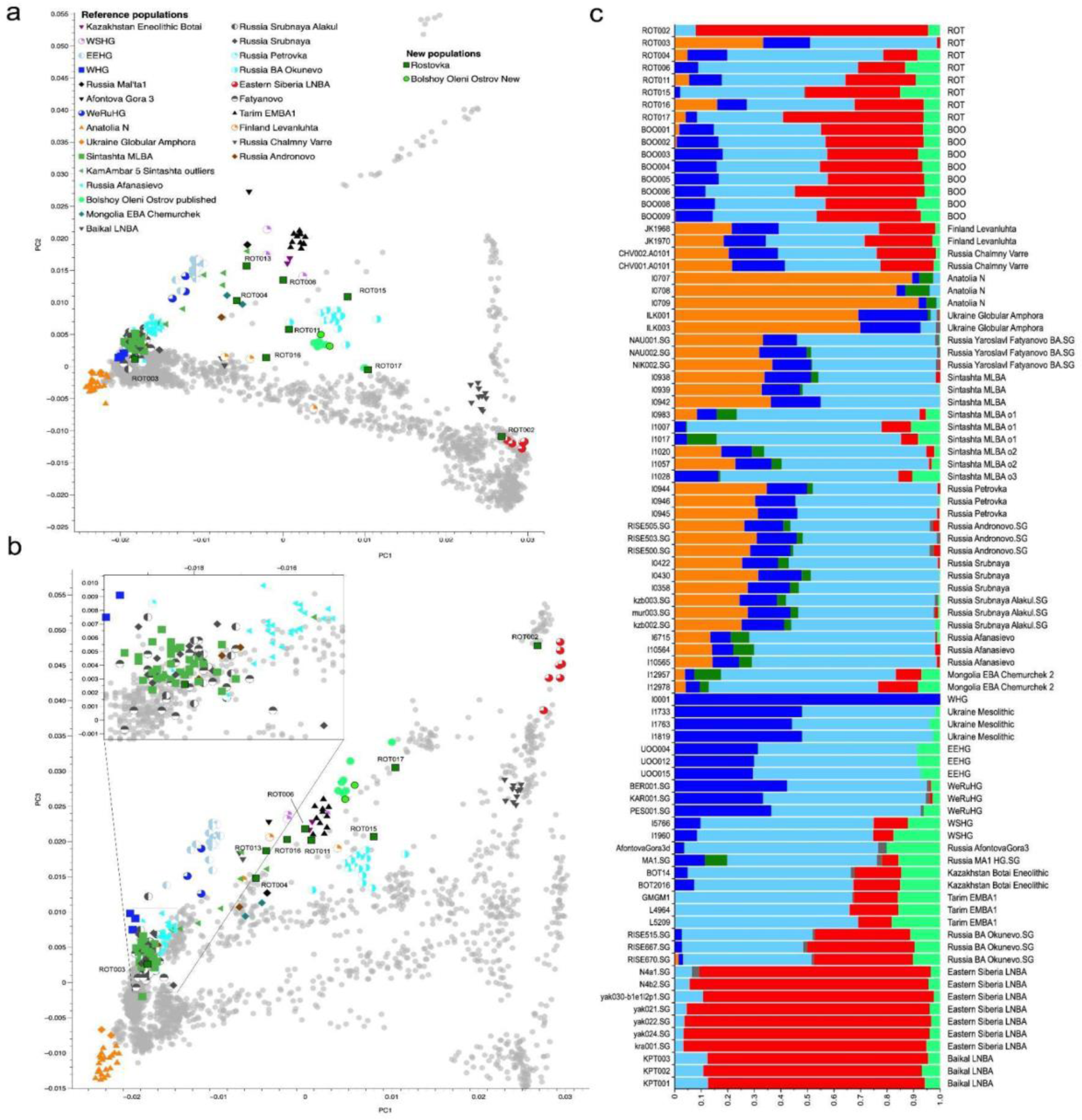
(a) PCA results with ancient individuals projected onto modern variation calculated using modern Eurasian populations. Modern samples are shown in gray. Ancient reference individuals are listed under “Reference populations”, and the new individuals are listed under “New populations”. PC1 vs PC2 are plotted; (b) PCA results for PC1 vs PC3; (c) Unsupervised ADMIXTURE results with the relevant populations and sample names shown (k=7).

### F-statistics and qpAdm

We calculated various F-statistics^29^ to formally assess the relationship of the ROT and BOO individuals with each other, and with different modern and ancient reference individuals and populations. First, we performed outgroup f3-statistics of the form *f*_3_ (Mbuti; test, modern) to test for the affinity of each ROT and BOO individual with modern Eurasian populations (Supplementary Figure 6, Supplementary Table 2). The *f*_3_-statistics results mirrored the distribution of the samples in the PCA and ADMIXTURE analyses, wherein the individuals with higher proportions of Eastern_Siberia_LNBA ancestry (e.g. ROT002) showed a greater affinity to the modern-day Siberian and Uralic-speaking populations, such as Nganasan, Evenk, Negidal, Nanai, and Ulchi (Supplementary Figure 6A), whereas the individuals with more Sintashta-like Western_Steppe_MLBA ancestry (e.g. ROT003) were closer to modern-day (North) Europeans, including Norwegian, Belarusian, Lithuanian, Scottish and Icelandic individuals (Supplementary Figure 6B). Comparisons with ancient groups using *f*_3_(Mbuti; test, published ancient) showed a similar trend (Supplementary Figure 6). For example, ROT002 on the ‘eastern end’ of the Eurasian cline, had the highest observed *f*_3_-values, i.e. shared more genetic drift, with Eastern_Siberia_LNBA, Russia Ust Belaya Neolithic, and Mongolia Early Iron Age individuals (Supplementary Figure 6A). By contrast, ROT003, the ‘westernmost’ individual in the Eurasian PCA space, had the highest affinity with Lithuania early Middle Neolithic Narva, Russia Sintashta, Kazakhstan Georgievsky Middle Bronze Age, Russia Poltavka, and Serbia Mesolithic individuals (Supplementary Figure 6 B). Similar trends could be observed for the BOO, wherein the modern Uralic-speaking populations, such as Nganasan and Selkup, were among the models with the highest *f*_3_-statistics. Among the ancient *f*_3_ comparisons, the most closely related individuals to BOO were the Eastern European and West-Siberian hunter-gatherers (EEHG and WSHG), published BOO, and Botai Eneolithic individuals from Kazakhstan (Supplementary Figures 6J and K).

Based on the geographic location of the sites, the results of the outgroup *f*_3_-statistics, and the distribution of the ROT and BOO individuals on the PCA, we tested whether these individuals retained more local Ancient North Eurasian (ANE) ancestry compared to contemporaneous groups and individuals from the general region. To assess the genetic affinities of ROT and BOO to known populations from similar general geographic area, time period, and archaeological affiliation, we calculated *f*_4_-statistics of the form *f*_4_(X, *test*; WSHG, Mbuti) where X stood for ROT and BOO individuals, and *test* populations included Okunevo, Tarim_EMBA_1, Sintashta_MLBA, and Eastern_Siberia_LNBA (Fig. 3). This test allowed us to identify groups that are cladal with ROT and BOO (i.e., equidistantly related to WSHG), and cases wheere ROT and BOO may have additional affinity to ANE (represented here by WSHG from Tyumen oblast, Russia as the best spatial and temporal proxy). Based on the *f*_4_-statistics, we find that ROT and BOO individuals carry excess affinity to ANE when compared to Eastern_Siberia_LNBA (Fig. 3A) and Russia MLBA Sintashta (Fig. 3C), except for ROT002 and ROT003. All BOO individuals are symmetrically related with the Okunevo Bronze Age group indicating no additional affinity to ANE (Fig. 3B). However, we see more heterogeneity in ROT, with some individuals having significantly more, and others significantly less genetic affinity to WSHG compared to Okunevo (Fig. 3B). All but one individual (ROT013) have significantly less ANE ancestry compared to Tarim EMBA (Fig. 3D). The general observations from *f*_4_-statistics mirror the trends we see in the PCA, especially when PC1 is plotted against PC2, wherein ROT individuals vary with regards to their location on the ANE cline represented by Afontova Gora 3 and Mal’ta 1, while the BOO individuals are more homogeneous (Fig. 2B).

**Figure 3.**
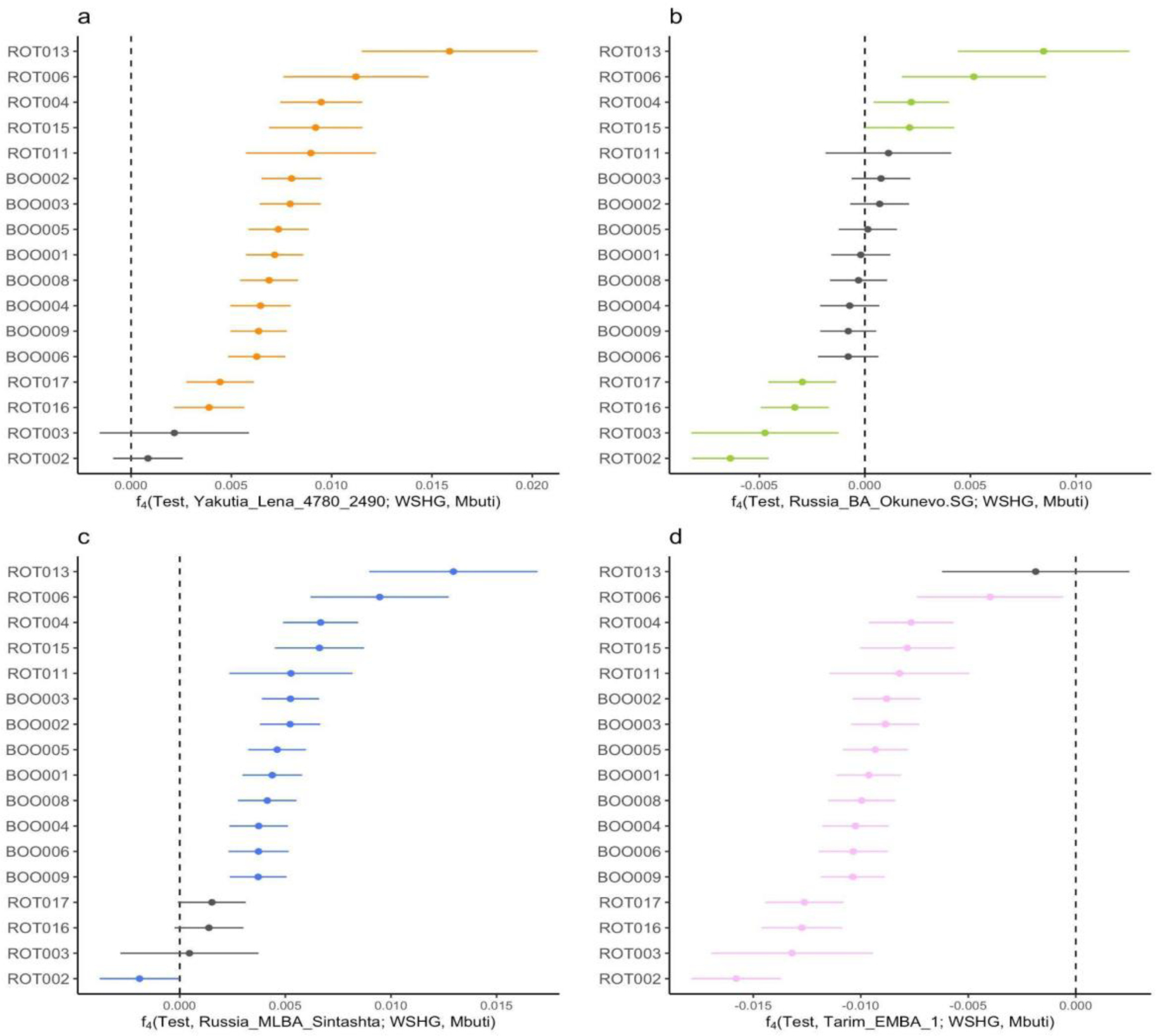
*f*_4_-statistics testing for excess ANE ancestry in ROT and BOO individuals. Testing for excess ANE ancestry with respect to: (a) Yakutia Lena 4780-2490, (b) Okunevo, (c) Russia MLBA Sintashta, (d) Tarim EMBA1. Significantly non-zero *f*_4_-statistics (|Z|>3) are shown in color, and non-significant *f*_4_-statistics are shown in gray. All error bars indicate 3 standard errors. “Test” denotes the individuals given on the y-axis.

The cultural affiliation of the BOO individuals remains poorly understood. Based on the archaeological information, such as the presence of ‘Waffle’ ware ceramics that are similar to the Neolithic pottery from Yakutia and Chukotka^16^, it was hypothesized that the BOO individuals represent a westward migration of Siberian populations along the forest-tundra and forest-steppe zones. However, potential contacts with Scandinavian archaeological cultures, such as Vardøy Ware, have been proposed for BOO^16^. To test this, we calculated pairwise *f*_3_-statistics with different ancient populations from Scandinavia, which separated the BOO individuals, as well the ROT from the rest of the populations ranging between Mesolithic to Medieval time periods (Supplementary Figure 7). Together, these results suggest a non-local genetic origin for the BOO individuals, and no substantial levels of early farmer ancestry, consistent with PCA and admixture analyses.

Lastly, we performed qpAdm analysis to formally test for and quantify the admixture proportions in ROT and BOO individuals that we had identified in the previous analyses (Fig. 4, Supplementary Table 3). Here, we could successfully model the ROT individuals as a mix of three ancestries, Eastern_Siberia_LNBA, Sintashta_MLBA, and WSHG, except for ROT002, which could be modeled instead as a two-source mixture of mainly Eastern_Siberia_LNBA ancestry and a smaller proportion of EEHG-like ancestry that could be represented by either Sintashta_MLBA, WSHG, or EEHG, whereas ROT003 could be modeled with Sintashta_MLBA as single source (Fig. 4B). We also tested whether ROT individuals could be modeled as a two-way mixture of the Eastern_Siberia_LNBA ancestry and either Sintashta_MLBA or WSHG as sources, however, this combination of ancestries did not result in consistently plausible model fits, compared to the combination of all three ancestries (Fig. 4a-c). By contrast, BOO individuals could not be modeled using either the combination of all three ancestry sources (Eastern_Siberia_LNBA, Sintashta_MLBA, and WSHG), or just a two-way mixture (Fig. 4a and c, Supplementary Table 3). However, replacing WSHG with EEHG as the putative local hunter-gatherer ancestry stratum and using Eastern_Siberia_LNBA as a second source provided good model fits (Fig. 4D, Supplementary Table 4).

**Figure 4.**
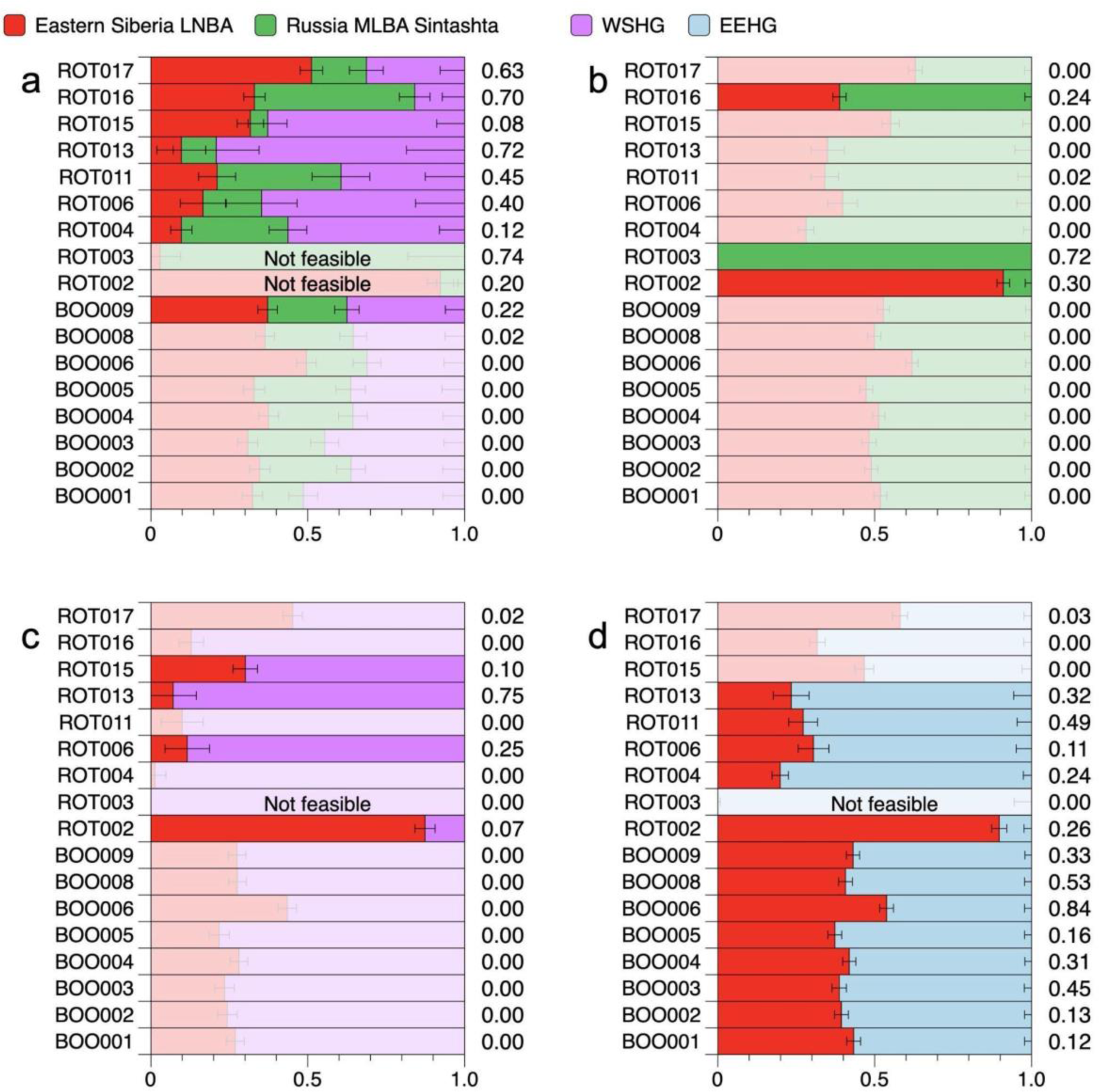
Ancestry modeling results for ROT and BOO individuals. (a) qpAdm models using Eastern Siberia LNBA, Russia MLBA Sintashta, and WSHG as sources; (b) qpAdm models with Eastern Siberia LNBA and Sintashta as sources; (c) qpAdm models with Eastern Siberia LNBA and WSHG as sources; (d) qpAdm models with Eastern Siberia LNBA and EEHG as sources. Corresponding p-values for each analysis are shown to the right of each row. Models with p-values < 0.05 are grayed out, and the models with negative ancestry proportions are indicated as “Not feasible”.

We then estimated the date of the admixture event in BOO individuals between the EEHG and Eastern_Siberia_LNBA sources using DATES v.753^30^. The admixture date was estimated to be 17.98±1.06 generations ago, or around 500 calendar years prior to the mean radiocarbon date of BOO, assuming a generation time of 29 years^31^ (Supplementary Figure 8).

### Identity-by-descent (IBD) analysis

We were interested in investigating distant biological relatedness among the BOO individuals (ROT individuals are below the required coverage threshold for imputation). To do so, we first imputed the genomes of the BOO individuals using GLIMPSE^32^ with the 1000G dataset^33^ as a reference panel. Based on the identification of haplotype blocks of certain lengths that are shared between individuals, i.e. identical by descent^34^, we confirmed the 2^nd^ degree related pair identified with READ (BOO004-BOO005), we also found two 3^rd^ degree related pairs (BOO003-BOO004 and BOO003-BOO005), as well as multiple potential 4^th^/5^th^-degree related pairs. The fact that the BOO individuals are distantly related to each other explains the relative homogeneity seen in the sample compared to ROT. With regards to the archaeological data from these individuals, two pairs of biologically related individuals were buried in the same graves, one 4^th^/5^th^-degree related pair: BOO005 (burial 17, sepulture 3, female) and BOO009 (burial 17, sepulture 4, female), and one 3^rd^-degree related pair: BOO003 (burial 16, sepulture 1, female) and BOO004 (burial 16, sepulture 3, male)^16^.

We also looked at IBD sharing between BOO and previously published individuals from groups that are broadly contemporaneous chronologically and close geographically, including Tarim_EMBA^35^, Okunevo^36^, Sintashta_MLBA^27^, EEHG^37^, Botai^36^, Yamnaya^36^, Easter_Siberia_LNBA^28^, and others (Fig. 5a, Supplementary Table 5). We found three shared IBD fragments (14-22cM) between BOO individuals and Sintashta_MLBA individuals (Supplementary Table 5), potentially suggesting shared ancestors as recent as approximately 500-750 years, and most likely reflecting the shared EEHG ancestry that is present in both groups.

**Figure 5.**
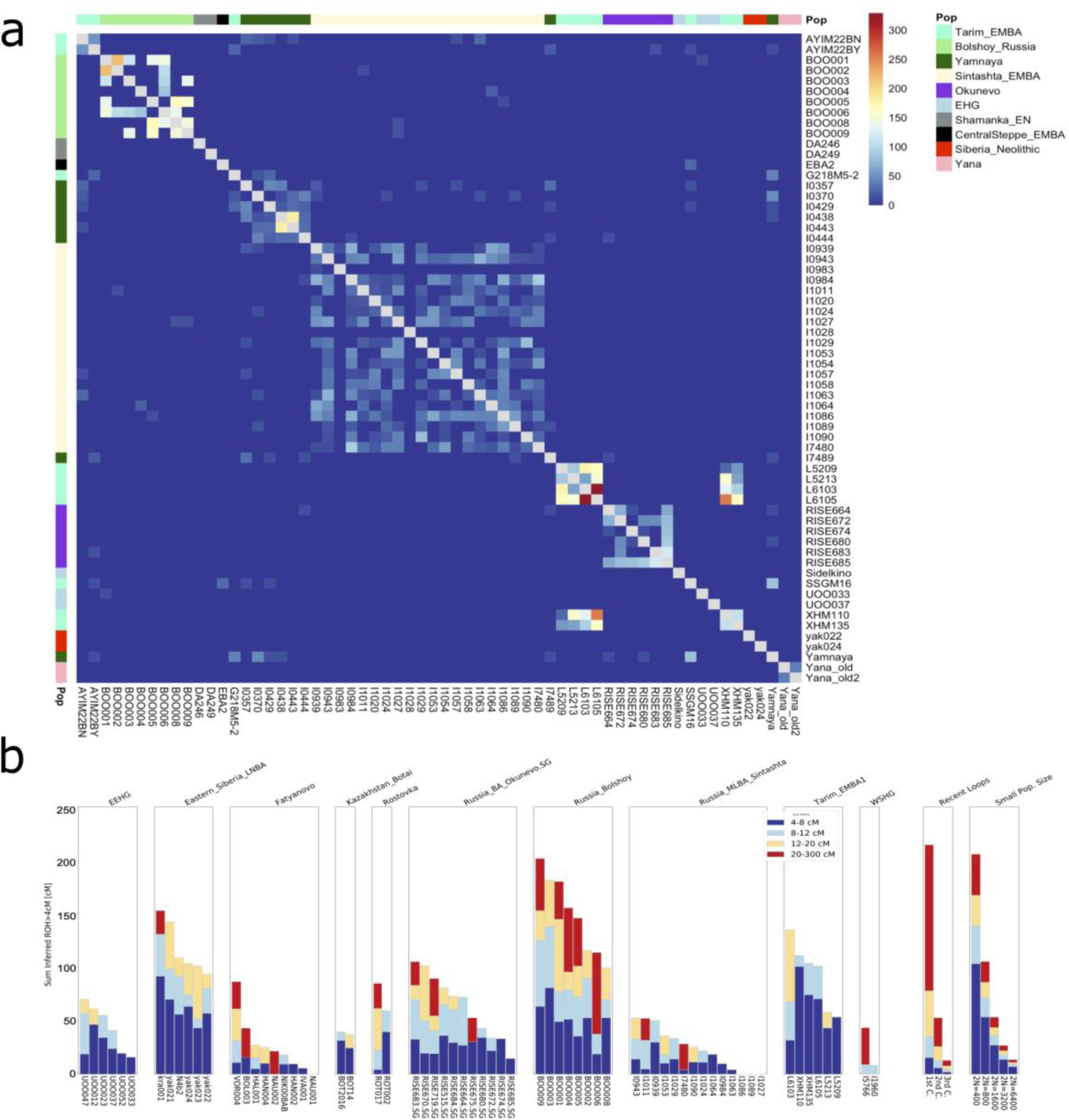
(a) IBD sharing between BOO and published data. Shared IBD chunks between 12 and 30 cM are shown. The total IBD length shared is indicated by the color of the square, and population designation is shown on the y-axis. (b) HapROH output for BOO, ROT and relevant contemporaneous populations. Runs of homozygosity (ROH) are plotted by population for individuals with more than 400k SNPs on the 1240k capture. ROH segments are colored according to their binned lengths.

Runs of homozygosity. To get a sense of the underlying population structure, general relatedness, and effective population sizes, we used HapROH to analyze runs-of-homozygosity (ROH) in the genomes of the BOO individuals, together with already published individuals with more than 400k SNPs on the 1240k SNP set^38^. We compared BOO to geographically and genetically close populations from the Eurasian forest steppe area, including Okunevo, Sintashta_MLBA, EEHG (UOO), Eastern_Siberia_LNBA, Tarim EMBA, and Fatyanovo (Fig. 5b). We also included two ROT individuals with more than 200K SNPs, but their results should be interpreted with caution. The ROH analysis of BOO suggests that this early Metal Age population had a relatively small effective population size of ∼2N=800, and one of the individuals (BOO006) appears to be an offspring of 2^nd^ cousins. Tarim EBMA, Okunevo, and Eastern_Siberia_LNBA groups also seemed to have relatively small effective population sizes, while Fatyanovo and Sintashta potentially had larger effective population sizes (Fig. 5b). In comparison, ROT individuals show similar ROH profiles to the populations they are closely related to, based on the PCA and F-statistics, i.e., ROT002 resembles the Eastern Siberian LNBA, and ROT017 – BOO (Fig. 5b).

### Demographic modeling

High-coverage shotgun data from BOO004 (∼30x) allowed us to perform demographic modeling to investigate North Eurasian genetic ancestry and the nature of the admixture of the Eastern and Western Eurasian sources found in BOO individuals using a site-frequency spectrum (SFS) modeling-based method called momi2^39^. We included published data from representative North Eurasian populations, both preceding and contemporaneous to BOO. After an incremental build-up of our model and including three admixture events, our final model indicates a recent admixture for BOO individuals (95% confidence interval (CI) 3596-4429 years ago), with substantial gene flow (39.9%; 95% CI 34.0-44.8%) from Eastern Eurasians (represented here by Late Neolithic/Bronze Age Siberian individuals), which is consistent with the results from qpAdm. The population size estimated for BOO (N=190, 95% CI 6-482) from momi2 (Fig. 6, Supplementary Table 6) is smaller than the estimate obtained from hapROH (2N between 400 and 800 individuals, Fig. 6). This could be explained by momi2 not taking into account inbreeding via the analysis of the runs of homozygosity, and thus producing a biased estimate of the true effective population size. Thus, we believe that the results produced by hapROH are closer to the true value of the effective population size.

**Figure 6.**
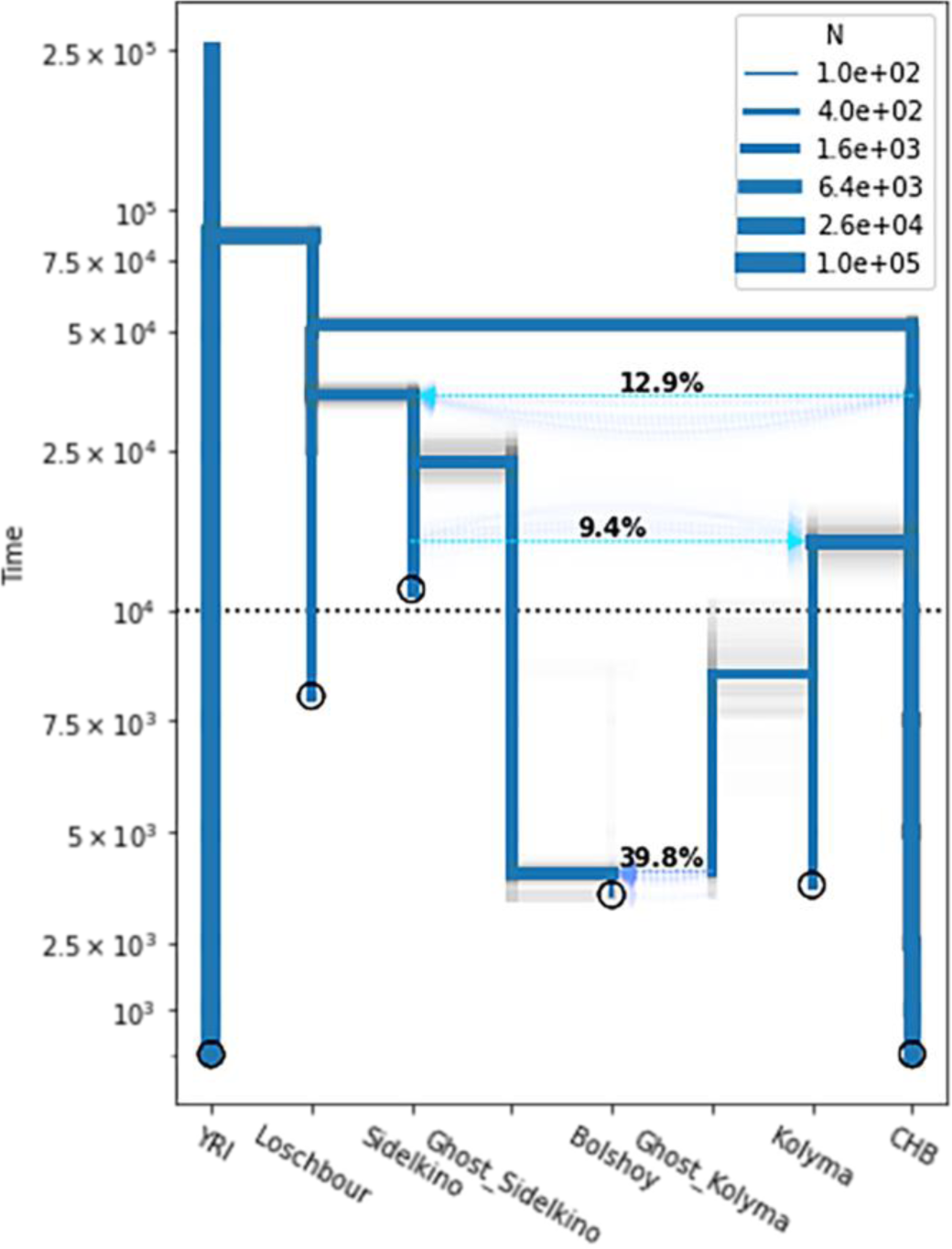
Momi2 demographic model for BOO004 using shotgun sequencing data from published ancient and modern individuals. Point estimates of the final model are shown in blue; results for 100 nonparametric bootstraps are shown in gray. The sampling times of populations are indicated by circles. The population sizes are indicated by the thickness of branches. The y-axis is linear below 10,000 years ago, and logarithmic above it. See Supplementary Table 6 for specific parameter values. YRI, Yoruba; CHB, Han Chinese.

### Y-chromosome haplogroups

We performed Y-haplogroup (Y-hg) typing of the ROT males using the YMCA method^19^ (Table 1). We identified two individuals that carried the R1a Y-hg (ROT003 (R1a-M417) and ROT016 (R1a-Z645)), one of the most widely distributed Y-hgs in Eurasia^40^. However, both individuals could be R1a-Z645, since ROT003 does not have either ancestral or derived ISOGG list SNPs after R1a-M417. The earliest evidence for Y-hg R1a comes from Mesolithic EEHG individuals^41,42^, and it is widespread throughout Eurasia today, from Scandinavia to South Asia and Siberia^40^. Specifically, R1a-M417 is common among Corded Ware-associated individuals^41,43^, while the derived R1a-Z645 is common in the Baltic Corded Ware^44^ and Fatyanovo^13^. Generally, due to their geographic distribution, these R1a haplogroups are thought to represent the eastward movement of the Corded Ware-, and Fatyanovo-associated groups.

ROT002, the individual with the highest proportion of north Siberian ancestry, was assigned to the N1a1a1a1a (N-L392) haplogroup. This Y-hg has also been found in two published BOO individuals^15^. The lineage N-L392 is one of the most common in present-day Uralic populations, and was estimated to have split from other N-lineages around 4,995 years ago^45^. This finding further highlights the importance of Y-hg N-L392 as being linked to the dissemination of proto-Uralic, but potentially also involving the ST network.

One of the males (ROT004) was assigned to haplogroup Q1b (Q-M346), which is found throughout Asia, including in several Turkic speaking populations, i.e. Tuvinians, Todjins, Altaians, Sojots, etc., and Mongolian-speaking Kalmyk population^46^. Another individual (ROT017) was determined to belong to the Q1b1 (Q-L53) haplogroup, which is common among present-day Turkic speakers across Eurasia. The branch Q-YP4004 includes Central Asian Q-L53(xL54) lineages and one ancient Native American individual from Lovelock Cave in Nevada dated 1.8 ky^47^, and the oldest individual with this haplogroup is irk040 (Cis-Baikal Neolithic, 4846 BP)^28^. The lineage C2a-L1373, carried by ROT011, is found at high frequency in Central Asian populations, North Asia and the Americas. C2a-L1373 expanded to North Asia and the Americas post-LGM, around 17,700–14,300 years ago^48^. Lastly, haplogroup R1b1a1a (R1b-M73), a sister-clade of R1b-M269, is carried by ROT006. This lineage is common in the Caucasus, Siberia, Mongolia, and Central Asia today^46^.

Overall, the Y-hg lineage diversity of male ROT individuals further highlights the heterogeneous nature of the ST, which has also been proposed by archaeologists^49^.

### Mitochondrial haplogroups

We identified a large diversity in the mitochondrial haplogroups (mt-hg) among ROT (Table 1), including mt-hgs that are found commonly in east Eurasia (A10, C1, C4, G2a1)^50–53^ and in west Eurasia (H1, H101, U5a, R1b, R1a)^36,54^, and similar to the trends seen from autosomal and Y-hg point of view. For example, the individual ROT002 with the highest affinity to Siberia_LNBA and carrying the Y-hg N-L392 also carries a mt-hg G2a1 commonly found in Eastern Eurasia. On the other hand, the individual ROT003 with the highest affinity to Sintashta_MLBA and carrying the Corded-ware derived Y-hg R1a1a1, is also a carrier of the R1a1a mt-hg commonly found in west Eurasia.

## Discussion

Metallurgical production is an important human cultural innovation that has been developed multiple times in multiple locations around the globe, one of which is Bronze Age Eurasia. It is during the Bronze Age period that the Seima-Turbino transcultural phenomenon is identified based on the evidence of skilled metallurgical production which is visible in the archaeological record of Northern Eurasia. The ST phenomenon holds an important place in the development of metallurgy within the framework of the Bronze Age Northern Eurasia, as the distribution of knowledge about the usage of Sn is one of its core elements^4^.

This study is reporting genome-wide data of ST-associated individuals and their connections to individuals associated with contemporaneous and preceding archaeological groups of the northern Eurasian forest-steppe and steppe during the Bronze Age, such as Sintashta, Okunevo, as well as Neolithic and Bronze Age Siberian groups. We also reassess the genetic structure of BOO individuals of northwestern Russia that have been shown to carry high levels of Siberian ancestry, an important characteristic of northern Eurasia.

The observed genetic heterogeneity among the ROT individuals can either suggest a group at an early stage of admixture, or signify the heterogeneous nature of the ST complex^4^. Together with evidence from the available archaeological data^4^, we argue that the individuals buried at ROT more likely represent a variety of genetic and perhaps cultural backgrounds, brought together by the ST metallurgical network. The findings from genome-wide autosomal data in PCA, ADMIXTURE and F-statistics are mirrored in the Y-chromosomal and mitochondrial data. Eight males of nine ROT individuals represent both eastern Eurasian and Western Eurasian Y-chromosomal lineages, and eastern and western Eurasian mitochondrial lineages, respectively, further highlighting the genetic heterogeneity seen in ROT. In general, the region of the Middle Irtysh around Rostovka can be characterized as the typological melting pot of the western and eastern part of the ST phenomenon mirroring in the genetic data.

On an individual level, there is no clear correlation between genetic ancestry of the screened individuals and the cultural/regional attribution of their grave goods. The inventory of the burial of individual ROT003 with western Eurasian genetic ancestry includes artifacts, which can be attributed to the eastern part of the ST phenomenon (forked lancehead with hook, trapezoidal socket axe) and there are regionally nonspecific ST-artifacts (lamellar dagger blades, forked lancehead with two loops), as well as ceramics of the Krotovo and Okunevo cultures. Artifacts with a connection to the Steppe-related cultures of the Transurals or the western part of the ST phenomenon are not present. The rather sparse burial of ROT002 does not stand out typologically in contrast to most other burials of the cemetery, despite the genetic attribution to the Eastern Siberian LNBA^4,55^.

In case of BOO, we see a very homogeneous genetic profile that we can model as a recent mixture of the Neolithic Siberian and EEHG components approximately ∼2200-2000 BCE, which places this event at a similar time as the temporal peak of the ST phenomenon. Interestingly, despite the geographic location of the burial site on the Kola Peninsula in northwestern Russia, BOO individuals carry higher proportions of ‘eastern’ Siberian ancestry than most ROT individuals. The genetic homogeneity observed in BOO individuals can be explained by the genetic background relatedness as shown by IBD sharing and ROH analysis, which is indicative of a relatively small or isolated population.

We also find that BOO and ROT exhibit distinct genetic subtleties with regard to the presence of the Early European Farmer ancestry. Although relatively contemporaneous, ROT individuals, in general, carry higher levels of Neolithic farmer-related ancestry, which we were able to model as part of the Sintashta_MLBA in our admixture models. However, this ancestry is not present in BOO individuals, which carry HG-related ancestry that is more similar to a more ancient, but local EEHG stratum (as demonstrated for the nearby Yuzhny Oleni Ostrov site)^41,56^. The lack of European farmer ancestry in BOO, contrary to what has been reported in Lamnidis 2018 (Figure 4a), also highlights the natural limits of the farming subsistence practice and the spread of farmer-related ancestry mediated by MBA forest steppe pastoralists into the northernmost parts of Eurasia during this time period.

We tested the individuals in this study with regards to the presence of Ancient North Eurasian ancestry, also known as the Upper Paleolithic Siberian ancestry that was first described in individuals from Mal’ta and Afontova Gora 2 and 3^56,57^, and marks a basal North Eurasian lineage that is shared between modern-day Europeans and Native Americans, but not found in southern India, East and Southeast Asia. This ancestry is generally associated with groups falling on the forest tundra genetic cline^25^, and is present in high levels in the Bronze Age Tarim mummies^35^. We found that the ST individuals vary with regards to their location on the ANE cline towards Afontova Gora 3 and Mal’ta 1, in line with the findings from PCA and ADMIXTURE analyses. BOO individuals also carry the ANE ancestry, but in a more homogeneous fashion.

With the new data from ROT, we are able to assess a recent proposal which suggested that Uralic languages could have been used within the ST network leading to the initial spread of Uralic languages across the Eurasian forest steppe^58–60^. After performing various tests of genetic structure of the ST-associated individuals, we report genetic profiles on an ancestry cline that generally mirrors the genetic distribution of modern-day Uralic-speaking populations of the northernmost forest-tundra (taiga and tundra) ecological cline^25^. Our findings show that the ST-associated individuals from Rostovka likely did not originate from a single location but rather represent people from a wide geographical area. Seima-Turbino was a latitudinal phenomenon on the same east-west axis where also the hypothetical homelands of the ancestral Uralic subgroups were positioned^61^. Thus, our genetic results are temporally and geographically consistent with the proposal that Uralic languages could have spread within the ST network, but are neither a clear nor a direct proof. Further ancient human DNA data from northern Eurasia will help elucidate the details of the wider spread of ancient Siberian ancestry and its association with proto-Uralic speaking groups.

Taken together, our findings show that all but one of the carriers of artifacts associated with the ST transcultural phenomenon have genetic similarities to the current taiga-tundra area populations, but harbor an extremely diverse mix of western and eastern Eurasian ancestries. However, due to the limited number of individuals studied, we cannot be certain as to what degree the individuals in this study represent the ST phenomenon as a whole. Genetic data from other confidently ST-associated sites will be crucial in providing a comparative analysis of the data. Lastly, we investigate the genetic history of the Siberian ancestry in northern Eurasia, and suggest that there were possibly several waves of migration of people carrying the Siberian ancestry component, indicating a complex demographic history of the region.

## Materials and Methods

### Archaeological background

Rostovka (ROT) is a ST burial site located on the river Om, in the city of Omsk, and was excavated in 1966-1969^9^. A total of 38 graves were excavated at the site, not all of which contained human remains. Some of the individuals buried at Rostovka were cremated, including ten clear cases of cremation, two graves with charcoal only, and one case with charcoal and the remains of children^55^.

The burial ground of Rostovka occupies a very curious position within the ST phenomenon. It is the largest ST-site east of the foothills of the Ural Mountains. It is, compared to other important sites like Seima and Turbino, well excavated, documented and published. Rostovka is, together with the smaller sites of Kargat 6 and Preobrazhenka 6, the most eastern point of the distribution of steppe-related artifacts in ST-contexts. At these three sites also specifically western ST-artifacts still appear, but in very small numbers. The ceramics of Rostovka are connected to the Krotovo and other forest-steppe cultures. The stone-artifacts are partly connected to the traditions of Baikalian stone-tools (little arrowheads, rectangular blades) partly to the western part of the ST phenomenon (bigger arrowheads, irregular blades) and in the case of a stone mace head to the Steppe-related cultures of the Transurals^55^. The metal composition of the artifacts of Rostovka is very comparable to the overall tendencies of the ST phenomenon. Steppe-related artifacts tend to contain less Sn than regional nonspecific ST-artifacts. And these nonspecific ST-artifacts contain less Sn than eastern ST-artifacts.

A total of 19 individuals from Rostovka were screened for ancient DNA preservation using shotgun sequencing of 5M reads, however, only nine passed the 0.1% endogenous DNA cutoff to be further analyzed using capture arrays. The low success rate is explained by the fact that the macroscopic preservation of the skeletal remains was poor in general, and we could only sample random parts of long bones and few teeth, but no petrous bones BOO was first excavated in 1925, with the most recent excavation taking place between 2001-2004^16^. A total of N=43 individuals were found, along with wooden grave constructions, as well as bone, antler, stone, ceramic, and bronze items. Most burials were inhumations, with the exception of three cremations, and most individuals were buried in wooden boat-shaped caskets^16^.

### DNA extraction and data generation

All aDNA work was done in dedicated clean laboratory facilities following the standard protocols^62^. Briefly, single-stranded libraries were produced for ROT, and double-stranded UDG-half libraries were produced for the new BOO individuals. First, shotgun libraries were screened for the presence of endogenous DNA, and samples with the aDNA content above 0.1% were captured for the 1240k sites. We also produced mtDNA and Y-haplogroup capture data for the samples included in the study. A set of BOO individuals were shotgun sequenced to high coverage. The nfcore/eager pipeline v.2.3.5^63^ was used to process the samples from fastq files to the deduplicated bam files. The software version information is listed in Supplementary Table 7. Briefly, samples were mapped to the hs37d5 version of the human reference genome using bwa aln with the following parameters: bwa aln -o 3 -n 0.001 -l 16500. Pseudohaploid genotyping calls for the ROT individuals were produced using pileupcaller (https://github.com/stschiff/sequenceTools) with the --singlestrandmode option. We trimmed two base pairs from bam files of BOO individuals from each side of the read, and genotyped the samples to produce pseudohaploid calls with pileupcaller (https://github.com/stschiff/sequenceTools). The ancient DNA status of the samples was authenticated using MapDamage v2 ^64^. Contamination from modern sources was determined using a combination of contammix^65^, schmutzi^66^, ANGSD X-chromosome contamination estimate (for males)^67^, and sex determination. READ^20^ and pairwise mismatch rate (PMR) were used to perform biological relatedness analysis. PMRs were calculated from pseudohaploid genotypes of the 1240k panel.

### Population genetics analyses

The projection PCA was done using smartpca^23^ including already published ancient and modern data from the Allen Ancient DNA Resource (AADR) v44.3^68^ using the projection mode, wherein ancient samples were projected upon modern genetic variation. Unsupervised admixture analysis was done on the ROT and the new BOO data together with already published ancient DNA samples from the AADR v44.3^68^ using ADMIXTURE^26^ for 1-20 K clusters between in 5 iterations. Coefficients of variance for each K were compared and the best K level was chosen based on the lowest average CV.

The f-statistics and qpAdm analyses were performed using admixr^69^. The resulting data were plotted using DataGraph v.4.6.1, and R^70^ using the ggplot2 package^71^. For qpAdm, we used Mbuti, Georgia_Kotias.SG, Israel_Natufian_published, Ami, Mixe, Italy_North_Villabruna_HG, and ONG.SG as an outgroup set (based on^15^).

Mitochondrial haplogroups were determined using HaploGrep2^72^ using the data from the mitochondrial capture. Briefly, mitochondrial capture data was mapped to the mitochondrial reference genome NC_012920.1 using circularmapper^73^ and mapping quality threshold of 30. Bam files were then imported into Geneios and a consensus fasta file was produced with the coverage threshold of 5, and Sanger heterozygotes set to >50%. The consensus fasta file was then imported into HaploGrep2. Y-haplogroup data generated using YMCA was used to assign Y-chromosome haplogroups to male ROT individuals following the method described in^19^.

ROH analysis was done using HapROH^38^ on the pseudohaploid data from BOO, together with already published individuals, and only focusing on samples with more than 400k SNPs from the 1240k SNP array.

BOO samples were imputed and phased using GLIMPSE^32^ following the default parameters, and merged with already published data, in order to test for patterns of IBD sharing among the individuals using ancIBD^34^. IBD analyses were restricted to samples covering more than 600K SNPs with GP>=0.99 after genotype imputation. IBD results were plotted using the R package pheatmap^74^.

### Demographic modeling

We used DATES^30^ to determine the time of admixture in BOO using Yakutia_Lena and UOO as the two reference sources. Demographic modeling of BOO was then performed using momi2^39^. To do so, we added several already published samples into our model to give sufficient background information: two random individuals from YRI (Yoruba in Ibadan, Nigeria) in 1000 Genomes Project^33^ representing the Africans; two random individuals from CHB (Han Chinese in Beijing, China) in 1000 Genomes Project representing East Asians; the 8000-year-old Loschbour individual from Luxembourg^75^ representing WEHG; one Mesolithic individual from Sidelkino, Russia^76^ representing EEHG; two Late Neolithic/Bronze Age individuals from Kolyma river regions of Yakutia, Russia ^28^ representing the Eastern Siberia LNBA. Ancient samples were downloaded from the European Nucleotide Archive (ENA) as bam files, and then processed and imputed using GLIMPSE, following the default parameters^32^. Modern individuals were extracted from the 1000G database^33^. We assumed a mutation rate of 1.25×10^-8^ per site per generation^77^ and a generation time of 29 years^31^. We progressively added more populations into the model. For each step we randomly initialized the new parameters dozens of times and did optimization respectively, and selected the best model as the basis of the next step. The initial model involves Western Eurasians (Loschbour) and Eastern Eurasians (CHB) as a simple split, with Africans (YRI) the outgroup. Each lineage was modeled to have its own population size. We also defined the ancestral Eurasian population size and the ancestral modern human size, and let the size of Loschbour and CHB exponentially change from the ancestral Eurasian size. We then added the EEHG (Sidelkino) lineage onto Loschbour, and Eastern Siberia LNBA (Kolyma) onto CHB, with their own population sizes. The shared ANE ancestry in Sidelkino and Kolyma was modeled as gene flow between Western and Eastern Eurasians. In actual modeling, we defined gene flow from ancestral Eastern Eurasians to Sidelkino, as well as gene flow from Sidelkino to Kolyma. At last, we added the highest coverage BOO individual BOO004 (Bolshoy) into the model, as an admixture of Sidelkino and Kolyma. We finally defined the ghost lineage for two source populations, with the same population size as their original branch, and modeled the Bolshoy lineage as the admixture of ghost lineages. When optimizing the final model, we got a series of similar likelihood results with recent admixture time and small population size in Bolshoy Oleni Ostrov lineage. We chose the final model whose admixture time matches the conclusion in DATES. To examine the stability of the parameters, we conducted 100 nonparametric bootstraps, fit the model for each bootstrap dataset using the parameter values of the final model as initial values, and computed the 95% confidence interval for each parameter. Each bootstrap dataset was created by dividing the whole genome into 100 equal-sized blocks and resampling the same number of blocks with replacement.

## Data Availability

Genomic data (BAM and fastq formats) are available on the European Nucleotide Archive (ENA) under accession number PRJEBXXX, genotypes in eigenstrat format can be found at https://edmond.mpdl.mpg.de.

## Author Contributions

Conceptualization: WH, AC Methodology: LS, FA Investigation: AC, WH, FF, ABR, LH Visualization: AC, WH, FF Supervision: WH, JK, SS Resources: FF, VM, VK, IK, SK Writing—original draft: AC, WH Writing—review & editing: WH, AC, ABR, FF, LH, SK, OV

## Classification

Social sciences, Anthropology, Population Genetics.

## Acknowledgments

We would like to thank all members of the Department of Archaeogenetics at the Max Planck Institute for Evolutionary Anthropology, the PALEoRIDER & Population Genetics Group. We also thank Dr. Elina Salmela for her suggestions and comments, and Dr. Rüdiger Krause. This work was funded by the European Research Council (ERC) under the European Union’s Horizon 2020 research and innovation program under grant agreement no, 771234-PALEoRIDER (to W.H.). Kuzminykh S.V. was supported by the program from the Archaeology Institute of the Russian Academy of Sciences No. NIOKTR 122022200264-9. Fabian Fricke was funded through a DFG grant titled “Zur Metallurgie der bronzezeitlichen Artefakte des Fundplatzes Sajtanska (nördlich von Ekaterinburg) vom Typ Sejmo-Turbino in Eurasien” (Dr. Rüdiger Krause).

## References

1. Chernykh, E. N. Ancient Metallurgy in the USSR: The Early Metal Age. (CUP Archive, 1992).

2. Kohl, P. L. The Making of Bronze Age Eurasia. (Cambridge University Press, 2007).

3. Hanks, B. K., Epimakhov, A. V. & Renfrew, A. C. Towards a refined chronology for the Bronze Age of the southern Urals, Russia. Antiquity 81, 353–367 (2007).

4. Chernykh, E. N. & Kuzminykh, S. V. Древняя металлургия Северной Евразии (сейминско-турбинский феномен). https://elibrary.ru/item.asp?id=21143678 (1989).

5. Linduff, K. M. Metallurgy in ancient eastern Eurasia. in Encyclopaedia of the History of Science, Technology, and Medicine in Non-Western Cultures 3103–3116 (Springer Netherlands, 2016).

6. Кузьминых, С. В. Сейминско-турбинская проблема: новые материалы. Краткие сообщения Института археологии 240–263 (2011).

7. Korochkova, O. Sacred Place of the First Metallurgists in the Middle Ural. (Издательство Уральского университета, 2020).

8. Parpola, A. Formation of the Indo-European and Uralic (Finno-Ugric) language families in the light of archaeology: Revised and integrated “total” correlations. https://researchportal.helsinki.fi/files/127256289/Parpola_A_2012._Formation_of_the_Indo_European_and_Uralic_language_families_in_the_light_of_archaeology._MSFOu_266.pdf (2012).

9. Marchenko, Z. V., Svyatko, S. V., Molodin, V. I., Grishin, A. E. & Rykun, M. P. Radiocarbon Chronology of Complexes With Seima-Turbino Type Objects (Bronze Age) in Southwestern Siberia. Radiocarbon 59, 1381–1397 (2017).

10. Chernykh, E. N. et al. Issues in the calendar chronology of the seima-turbino transcultural phenomenon. Archaeol. Ethnol. Anthropol. Eurasia (Russ.-lang.) 45, 45–55 (2017).

11. Ковтун, И. В., Марочкин, А. Г. & Герман, П. В. Радиоуглеродные даты и относительная хронология сейминско-турбинских, крохалёвских и самусьских древностей. Труды V (XXI) Всероссийского (2017).

12. Tambets, K. et al. Genes reveal traces of common recent demographic history for most of the Uralic-speaking populations. Genome Biology vol. 19 Preprint at 10.1186/s13059-018-1522-1 (2018).

13. Saag, L. et al. The Arrival of Siberian Ancestry Connecting the Eastern Baltic to Uralic Speakers further East. Curr. Biol. 29, 1701–1711.e16 (2019).

14. Der Sarkissian, C., et al. Ancient DNA reveals prehistoric gene-flow from siberia in the complex human population history of North East Europe. PLoS Genet. 9, e1003296 (2013).

15. Lamnidis, T. C. et al. Ancient Fennoscandian genomes reveal origin and spread of Siberian ancestry in Europe. Nat. Commun. 9, 5018 (2018).

16. Murashkin, Kolpakov & Shumkin. Kola Oleneostrovskiy grave field: a unique burial site in the European Arctic. Iskos (2016).

17. Mathieson, I. et al. Genome-wide patterns of selection in 230 ancient Eurasians. Nature 528, 499–503 (2015).

18. Fu, Q. et al. An early modern human from Romania with a recent Neanderthal ancestor. Nature 524, 216–219 (2015).

19. Rohrlach, A. B. et al. Using Y-chromosome capture enrichment to resolve haplogroup H2 shows new evidence for a two-path Neolithic expansion to Western Europe. Sci. Rep. 11, 15005 (2021).

20. Monroy Kuhn, J. M., Jakobsson, M. & Günther, T. Estimating genetic kin relationships in prehistoric populations. PLoS One 13, e0195491 (2018).

21. Lipatov, M., Sanjeev, K., Patro, R. & Veeramah, K. R. Maximum Likelihood Estimation of Biological Relatedness from Low Coverage Sequencing Data. bioRxiv 023374 (2015) doi:10.1101/023374.

22. Ramsey, C. B. Bayesian Analysis of Radiocarbon Dates. Radiocarbon 51, 337–360 (2009).

23. Patterson, N., Price, A. L. & Reich, D. Population structure and eigenanalysis. PLoS Genet. 2, e190 (2006).

24. Wang, C.-C. et al. Ancient human genome-wide data from a 3000-year interval in the Caucasus corresponds with eco-geographic regions. Nat. Commun. 10, 1–13 (2019).

25. Jeong, C. et al. The genetic history of admixture across inner Eurasia. Nat Ecol Evol 3, 966–976 (2019).

26. Alexander, D. H., Novembre, J. & Lange, K. Fast model-based estimation of ancestry in unrelated individuals. Genome Res. 19, 1655–1664 (2009).

27. Narasimhan, V. M. et al. The formation of human populations in South and Central Asia. Science 365, (2019).

28. Kılınç, G. M. et al. Human population dynamics and Yersinia pestis in ancient northeast Asia. Sci Adv 7, (2021).

29. Patterson, N. et al. Ancient admixture in human history. Genetics 192, 1065–1093 (2012).

30. Chintalapati, M., Patterson, N. & Moorjani, P. Reconstructing the spatiotemporal patterns of admixture during the European Holocene using a novel genomic dating method. bioRxiv 2022.01.18.476710 (2022) doi:10.1101/2022.01.18.476710.

31. Fenner, J. N. Cross-cultural estimation of the human generation interval for use in genetics-based population divergence studies. Am. J. Phys. Anthropol. 128, 415–423 (2005).

32. Rubinacci, S., Ribeiro, D. M., Hofmeister, R. J. & Delaneau, O. Efficient phasing and imputation of low-coverage sequencing data using large reference panels. Nat. Genet. 53, 120–126 (2021).

33. 1000 Genomes Project Consortium et al. A global reference for human genetic variation. Nature 526, 68–74 (2015).

34. Ringbauer, H. et al. ancIBD -Screening for identity by descent segments in human ancient DNA. *bioRxiv* 2023.03.08.531671 (2023) doi:10.1101/2023.03.08.531671.

35. Zhang, F. et al. The genomic origins of the Bronze Age Tarim Basin mummies. Nature 599, 256–261 (2021).

36. de Barros Damgaard, P., et al. The first horse herders and the impact of early Bronze Age steppe expansions into Asia. Science 360, eaar7711 (2018).

37. Posth, C. et al. Palaeogenomics of upper Palaeolithic to neolithic European hunter-gatherers. Nature 615, 117–126 (2023).

38. Ringbauer, H., Novembre, J. & Steinrücken, M. Parental relatedness through time revealed by runs of homozygosity in ancient DNA. Nat. Commun. 12, 1–11 (2021).

39. Kamm, J., Terhorst, J., Durbin, R. & Song, Y. S. Efficiently inferring the demographic history of many populations with allele count data. J. Am. Stat. Assoc. 115, 1472–1487 (2020).

40. Underhill, P. A. et al. The phylogenetic and geographic structure of Y-chromosome haplogroup R1a. Eur. J. Hum. Genet. 23, 124–131 (2015).

41. Haak, W. et al. Massive migration from the steppe was a source for Indo-European languages in Europe. Nature 522, 207–211 (2015).

42. Saag, L. et al. Genetic ancestry changes in Stone to Bronze Age transition in the East European plain. Sci Adv 7, (2021).

43. Papac, L. et al. Dynamic changes in genomic and social structures in third millennium BCE central Europe. Sci Adv 7, (2021).

44. Saag, L. et al. Extensive Farming in Estonia Started through a Sex-Biased Migration from the Steppe. Curr. Biol. 27, 2185–2193.e6 (2017).

45. Ilumäe, A.-M. et al. Human Y Chromosome Haplogroup N: A Non-trivial Time-Resolved Phylogeography that Cuts across Language Families. Am. J. Hum. Genet. 99, 163–173 (2016).

46. Malyarchuk, B. et al. Ancient links between Siberians and Native Americans revealed by subtyping the Y chromosome haplogroup Q1a. J. Hum. Genet. 56, 583–588 (2011).

47. Grugni, V. et al. Analysis of the human Y-chromosome haplogroup Q characterizes ancient population movements in Eurasia and the Americas. BMC Biol. 17, 3 (2019).

48. Sun, J. et al. Post-last glacial maximum expansion of Y-chromosome haplogroup C2a-L1373 in northern Asia and its implications for the origin of Native Americans. Am. J. Phys. Anthropol. 174, 363–374 (2021).

49. Molodin & Durakov. The adaptation of the Seima-Turbino tradition to the Bronze Age cultures in the south of the West Siberian plain. *& Anthropology of …*.

50. Schurr, T. G., Sukernik, R. I., Starikovskaya, Y. B. & Wallace, D. C. Mitochondrial DNA variation in Koryaks and Itel’men: population replacement in the Okhotsk Sea-Bering Sea region during the Neolithic. Am. J. Phys. Anthropol. 108, 1–39 (1999).

51. Volodko, N. V. et al. Mitochondrial genome diversity in arctic Siberians, with particular reference to the evolutionary history of Beringia and Pleistocenic peopling of the Americas. Am. J. Hum. Genet. 82, 1084–1100 (2008).

52. Pilipenko, A. S., Trapezov, R. O., Zhuravlev, A. A., Molodin, V. I. & Romaschenko, A. G. MtDNA Haplogroup A10 Lineages in Bronze Age Samples Suggest That Ancient Autochthonous Human Groups Contributed to the Specificity of the Indigenous West Siberian Population. PLoS One 10, e0127182 (2015).

53. Tanaka, M. et al. Mitochondrial genome variation in eastern Asia and the peopling of Japan. Genome Res. 14, 1832–1850 (2004).

54. Ning, C. et al. Ancient Mitochondrial Genomes Reveal Extensive Genetic Influence of the Steppe Pastoralists in Western Xinjiang. Front. Genet. 12, 740167 (2021).

55. Matyushenko, V. I. & Sinitsina, G. V. Могильник у д. Ростовка вблизи Омска [Burial Ground near the Village of Rostovka near Omsk]. https://elibrary.ru/item.asp?id=24232508 (1988).

56. Fu, Q. et al. The genetic history of Ice Age Europe. Nature 534, 200–205 (2016).

57. Raghavan, M. et al. Upper Palaeolithic Siberian genome reveals dual ancestry of Native Americans. Nature 505, 87–91 (2014).

58. Asko, P. LOCATION OF THE URALIC PROTO-LANGUAGE IN THE KAMA RIVER VALLEY AND THE URALIC SPEAKERS’ EXPANSION EAST AND WEST WITH THE ’SEJMA-TURBINO TRANSCULTURAL PHENOMENON’ 2200-1900 BC. Археология евразийских степей 258–277 (2022).

59. Grünthal, R. et al. Drastic demographic events triggered the Uralic spread. Diachronica 39, 490–524 (2022).

60. Kovtun, I. V. Предыстория индоарийской мифологии [Prehistory of Indo-Aryan mythology]. (Азия-принт, 2013).

61. Saarikivi, J. The divergence of Proto-Uralic and its offspring. The Oxford Guide to the Uralic Languages 28–58 Preprint at 10.1093/oso/9780198767664.003.0002 (2022).

62. A Fellows Yates, J., et al. A-Z of ancient DNA protocols for shotgun Illumina Next Generation Sequencing v2. (2021) doi:10.17504/protocols.io.bvt9n6r6.

63. Fellows Yates, J. A., et al. Reproducible, portable, and efficient ancient genome reconstruction with nf-core/eager. PeerJ 9, e10947 (2021).

64. Jónsson, H., Ginolhac, A., Schubert, M., Johnson, P. L. F. & Orlando, L. mapDamage2.0: fast approximate Bayesian estimates of ancient DNA damage parameters. Bioinformatics 29, 1682–1684 (2013).

65. Fu, Q. et al. A revised timescale for human evolution based on ancient mitochondrial genomes. Curr. Biol. 23, 553–559 (2013).

66. Renaud, G., Slon, V., Duggan, A. T. & Kelso, J. Schmutzi: estimation of contamination and endogenous mitochondrial consensus calling for ancient DNA. Genome Biol. 16, 224 (2015).

67. Korneliussen, T. S., Albrechtsen, A. & Nielsen, R. ANGSD: Analysis of Next Generation Sequencing Data. BMC Bioinformatics 15, 356 (2014).

68. Mallick, S., et al. The Allen Ancient DNA Resource (AADR): A curated compendium of ancient human genomes. *bioRxiv* 2023.04.06.535797 (2023) doi:10.1101/2023.04.06.535797.

69. Petr, M., Vernot, B. & Kelso, J. admixr—R package for reproducible analyses using ADMIXTOOLS. Bioinformatics 35, 3194–3195 (2019).

70. R Core Team, A., Team, R. C. & Others. R: A language and environment for statistical computing. R Foundation for Statistical Computing, Vienna, Austria. 2012. Preprint at (2022).

71. Wickham, H. ggplot2: Elegant Graphics for Data Analysis. (Springer International Publishing, 2016).

72. Weissensteiner, H. et al. HaploGrep 2: mitochondrial haplogroup classification in the era of high-throughput sequencing. Nucleic Acids Res. 44, W58–63 (2016).

73. Peltzer, A. et al. EAGER: efficient ancient genome reconstruction. Genome Biol. 17, 60 (2016).

74. Kolde, R. pheatmap: Pretty heatmaps. (Github, 2012).

75. Lazaridis, I. et al. Ancient human genomes suggest three ancestral populations for present-day Europeans. Nature 513, 409–413 (2014).

76. Damgaard, P. de B., et al. 137 ancient human genomes from across the Eurasian steppes. Nature 557, 369–374 (2018).

77. Fu, Q. et al. Genome sequence of a 45,000-year-old modern human from western Siberia. Nature 514, 445–449 (2014).

