## Supplementary Information for "Bronze Age Northern Eurasian Genetics in the Context of Development of Metallurgy and Siberian Ancestry"

**This PDF file includes:**

Supplementary Text

Supplementary Figures 1-8

References

**Supplementary Note 1**

**Archaeological Background.** Generally, there is no drastic change in the metal complex, stone and bone inventory of the ST proper, however, some differences do exist. There are two traditions of manufacturing bronze celts: the Rostovka-Seima tradition (tools with and without hooks, decorated with a ladder belt and geometric figures - triangles and rhomboids), and the Turbino tradition (without side hooks, decorated with a belt of horizontal lines or without any ornamentation). Both traditions were continued, first of all, in the forest belt of Northern Eurasia in the post-Seima epoch until the end of the Early Iron Age.

Based on the analysis of ST artifacts (typology and metal alloy composition), a “western” (or European) and “eastern” (or Transural Siberian) variant of the phenomenon have been identified ^1^. For example, in the east of the Urals a greater number of casting molds have been identified compared to the west of the Urals. However, west of the Urals a greater number and variation of ST objects have been identified. Of note, burials of actual metal workers (blacksmiths) have only been found east of the Urals. The western ST can also be characterized by the presence of the so-called Eurasian component potentially associated with Abashevo cultures and others from the Southern Urals and the Volga-Kama River basin (Chernykh and Kuzminykh 1989). Geographic structure regarding the degree of similarity of typology among the ST objects has been shown, with the similarity decreasing with distance^1^.

Regarding their regional attribution, the typical artifacts of the ST phenomenon can be categorized into three groups. First, there are artifact types that can be found in the entire distribution area of the ST phenomenon. Those are the lamellar dagger blades of the types NK-2, NK-4 and NK-6, the socket axes of the types K-14, K-18, K-30, K-32 and K-34 and the forked lanceheads of the types KD-12, KD-14, KD-16, KD-18, KD-20, KD-22, KD-24 and KD-26. The lamellar dagger blades have in common a flat cross section without a mid-rib. The different types are mainly based on the size of the blades themselves. The different types of socket axes are all relatively heavily decorated with rhombuses, triangles and tram-line ornaments around the socket. The different types of forked lanceheads are mainly defined by the presence or absence of ornaments and loops on the socket^1^.

Special to the area from the Urals to the western end of the ST distribution area are the lamellar dagger blades of the types NK-6 and NK-8, the serrated dagger blades of the types NK-18, NK-20, NK-22 and NK-24, the socket axes of the types K-4, K-6, K-8, K-10, K-12 and K-16 and the fully hilted daggers of the types KZh-2, KZh-4, KZh-6 and KZh-8. The lamellar dagger blades from NK-6 and NK-8 distinguish themselves by a rhombic cross section with a slide mid-rib on the blade. Most of the specific socket axes of the western part of the STP are less ornamented with the expectation of K-16, which was exclusively found in Seima on the banks of the Volga. The fully hilted daggers of the types KZh-2 and KZh-4 show lamellar ST-style blades. The daggers from the types KZh-6 and KZh-8 have curved blades, which are closer in style to dagger blades of the steppe cultures (Sintashta, Petrovka, Alakul etc.), but fully hilted the nearly exclusively appear in ST-contextes. Beside these ST-specific types, artifacts of the aforementioned Steppe-related cultures also appear more often in the western part of the distribution area^1^.

The eastern part of the ST phenomenon east of the foothills of the Ural Mountains also provides specific kinds of objects. The types K-20, K-22, K-24, K-26 and K-28 distinguish themselves mostly by a heavier ornamentation and a more trapezoidal shape. The forked lanceheads of the types KD-6, KD-8 and KD-10 are extraordinarily large compared to the other types of these lanceheads. Examples of K-10 moreover show a large hook on the side of the socket. The knife blades of the type NK-28 are special, as they were found mounted in antler handles at 60°-90° degrees. The blade-style of the fully hilted knives of the types KZh-8 and KZh-10 is unique within the ST phenomenon and is most likely the base of the later development of the knives of the Siberian Karasuk culture ^1^.

To analyze the metallurgy of the aforementioned objects, we have to compare data that was obtained through very different methods during various periods of archaeometallurgic research. The oldest data stem from the very starting days of the Laboratory of Scientific Methods of the Academy of Science of the USSR from the early 1960s using Optically Stimulated Spectrometry. The newest data were obtained during the last decade in Russian and German laboratories operating on X-ray fluorescence. Because of that, we chose a rather robust approach, comparing just the most important alloying element for the metallurgy of the ST-era, tin (Sn).

We grouped all of the ST-sites in six different regional groups, following river systems and mountain ranges. The most western group is made up of the Baltic sites, however, there is no trace element analysis of this material. East of these sites, there is a concentration of ST-sites between the Dnipro and the Upper Volga. More to the east, there is a dense group of ST-related sites at the banks of the Middle and Lower Volga and the Kama. The agglomeration of ST-sites east of the Ural Mountains we describe as the Transural-Region. The area of the banks of the Ob and Irtysh rivers makes up another region, and finally the sites in the Altai and Tuva Mountains form the most eastward region (Supplementary Figure 1).

Going west to east, we observe a steady increase of Sn within ST-artifacts. The Dnipro/Upper Volga and Lower Volga/Kama regions are very comparable with a median of 2% Sn and 1% Sn and an upper end of the 1-sigma distribution around 6% Sn and lower end around 0% Sn each. The median of the Transural region is about 3% Sn and a slightly higher 1-sigma distribution of 1% to 7%. The objects of the Ob/Irtysh region though have a median of 8% Sn with a 1-sigma interval from 5% to 10% Sn. The Altai/Tuva region is even a bit higher with a median of 10% and a 1-sigma distribution from 7% to 11% Sn. The upper end of the 2-sigma interval for all regions is at about 15% Sn (Supplementary Figure 2).

The correlation between geographical location and the amount of Sn holds true when analyzed by typological groups (Supplementary Figure 3). For example, western-related socket ax types contain significantly less Sn compared to the eastern-related or nonspecific types. The difference between nonspecific and eastern-related types is smaller but is still recognisable. In the case of forked lanceheads, eastern-related types contain more Sn than nonspecific types (there are no specifically western-related types of forked lances). The ST-specific dagger blades are the exception to this rule. The few western-related dagger blades contain a higher amount of Sn than the unspecific ones (there are no eastern-related types of dagger blades), but they in general contain less Sn compared to forked lances or socket axes. These results once again confirm the theory that the Altai along with the other mountain ranges of Eastern Kazakhstan should be considered the source of Sn for the ST phenomenon.

Changes in the material complex of ST are connected with contacts and incorporations from other cultures. On the Irtysh River at the Rostovka site wrought spearheads and knives, typical for Sintashta and Petrovka cultures of the Trans-Urals steppes are found.

Turbino, Korshunovo, and Ust-Gaiva sites on the Middle Kama show the presence of axes, wrought spearheads, and knives. In Sokolovka at the mouth of the Kama River, in Ust-Vetluga at the mouth of the Vetluga River, in Seima and Reshnyi on the Oka River - axes, wrought spearheads, knives, and adornments typical for the Abashevo and Pokrovka (early Russian Srubnaya) antiquities are present.

Ceramics from the Siberian burial grounds of the ST (Rostovka and Satyga) show forms similar to the vessels of the Krotovo and other West Siberian cultures. In the Middle Kama burials and in Ust-Vetluga there are no ceramics in the burials. The vessels from Sokolovka, Seima and Reshnyi are Abashevo-like based on their shape (bell-shaped), ornamentation and alloy composition (shell).

**Rostovka site description.** At the level of burials of the screened individuals from Rostovka there are also several observations, which should be recognised. Out of the nice graves with successfully analyzed individuals, five contain ST-style metal artifacts. The grave of ROT013 contains no finds at all^2^.

The individual ROT002 was buried together with a dagger blade of the type NK-4, several nonspecific flint-tools, a grindstone, two ceramic vessels and a bone-arrowhead, similar to arrowheads, which were found among the ST-materials of the Kaminskaya Cave on the western site of the Urals^2^.

Grave 8, in which RO003 was buried, is particularly rich. It contains two lanceheads of the types KD-10 and KD-14, a socket ax of type K-20, a dagger blade of the type NK-6, several small flint arrowheads, two rectangular flint blades, and a bone handle^2^.

The grave goods, which were found together with individual ROT011 in grave 23 consist of a dagger blade of type NK-6, a rectangular flint blade, a piece of copper ore and a nonspecific polished stone tool^2^.

The inventory of grave 33, in which ROT016 was buried, contains a lancehead of type KD-24, two gold rings, several small flint arrowheads, two stone objects, which are usually described as arrow-smoothers, and a big number of perforated bone battens. These battens are interpreted as the remains of a piece of body-armor^2^.

ROT017 was buried in grave 34 with an inventory of a dagger blade of type NK-4, a socket ax of type 20, a lancehead of type KD-10, two gold rings, a big ceramic fragment, several small flint blades, and a little bone awl^2^.

ROT004 was buried together only with a little asymmetrical fint-blade and a piece of bone, which could be a fragment of a body-armor, like in grave 33. The grave of ROT006 contains only an asymmetrical flint-blade and a very little metal awl. ROT015 was found together with some little fragments of ceramics, a rectangular blade and several other flint-flakes^2^.

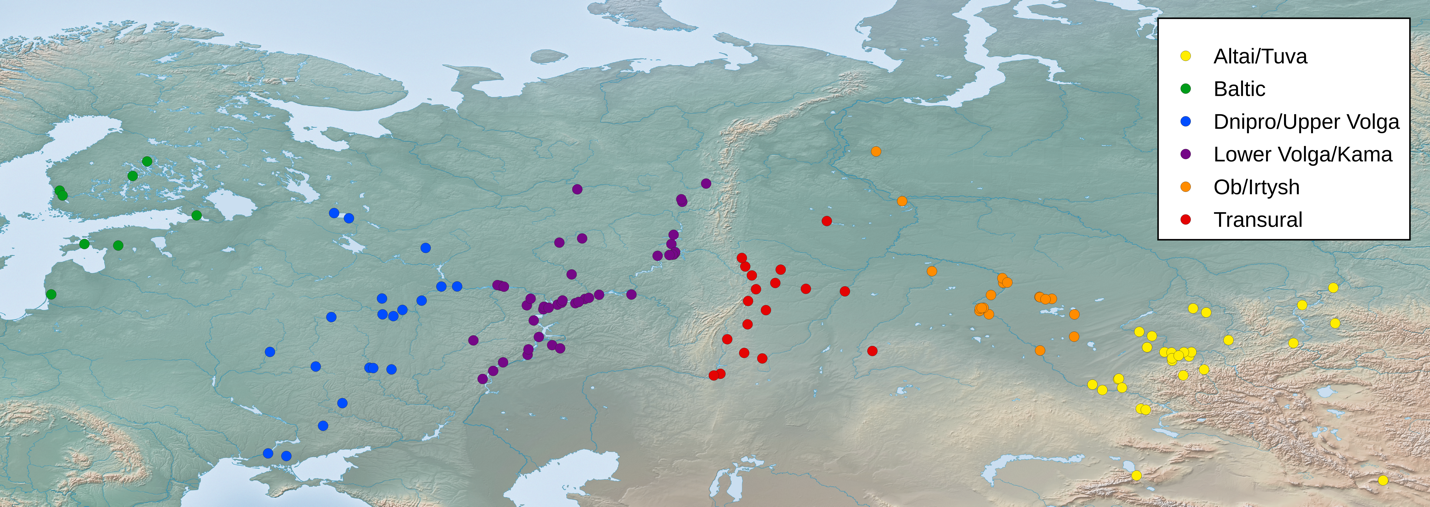

**Supplementary Figure 1.** Regional groupings of the sites of the STP

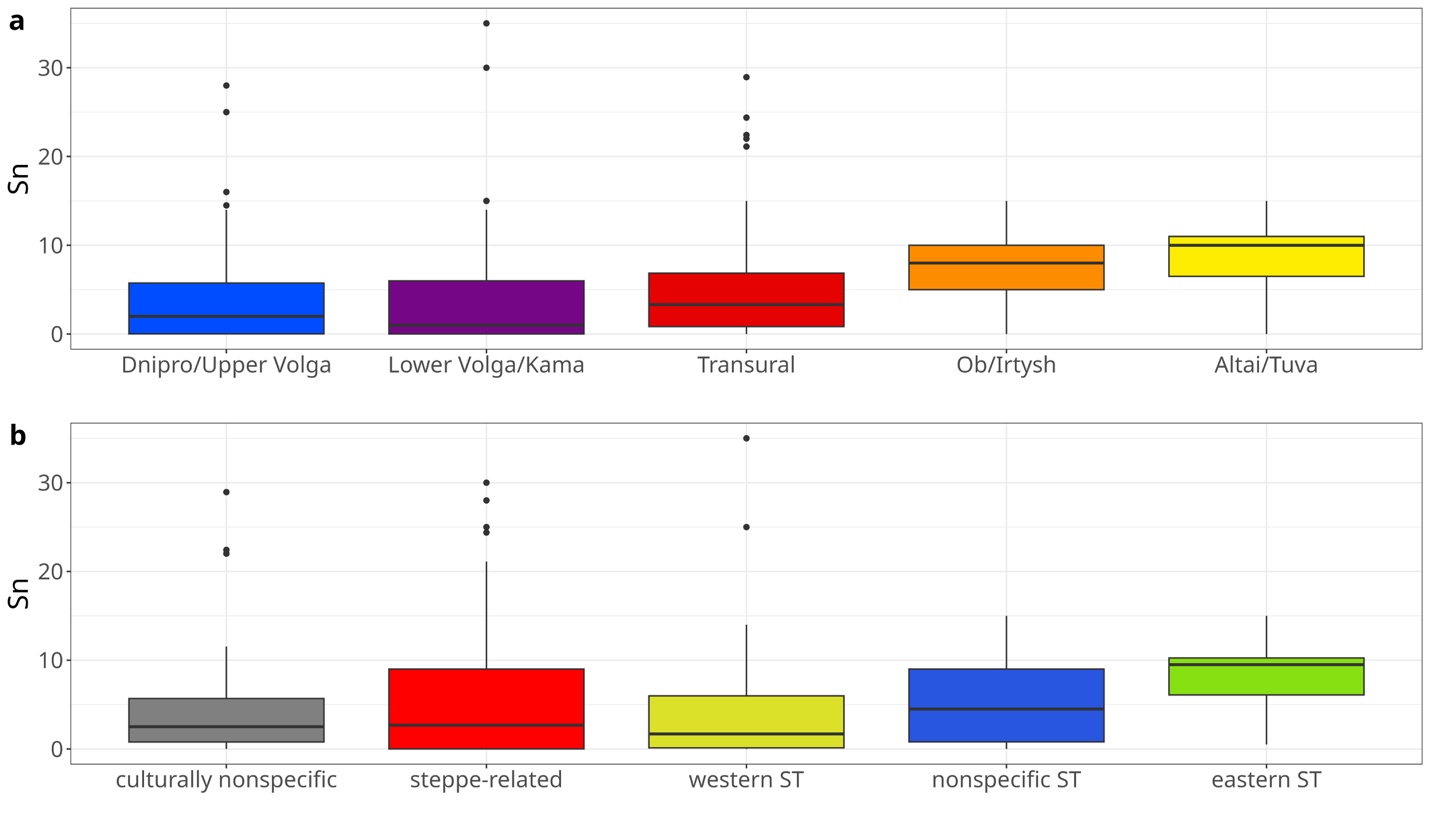

**Supplementary Figure 2.** Tin-content of the artifacts of ST-sites grouped by regions (a) and cultural attribution (b)

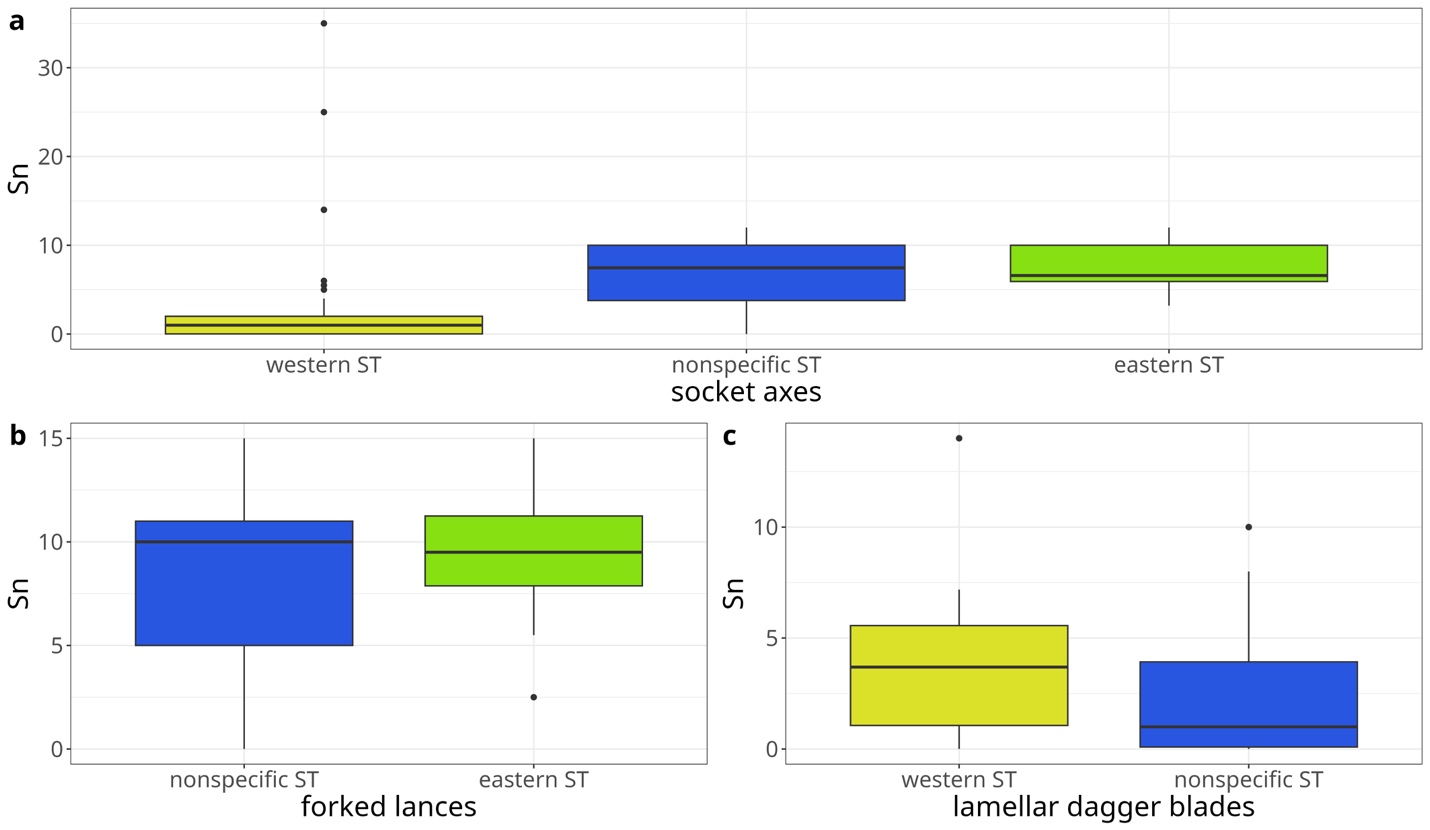

**Supplementary Figure 3.** Tin-content of socket axes (a), forked lances (b) and lamellar dagger blades (c) grouped by cultural attribution of their types.

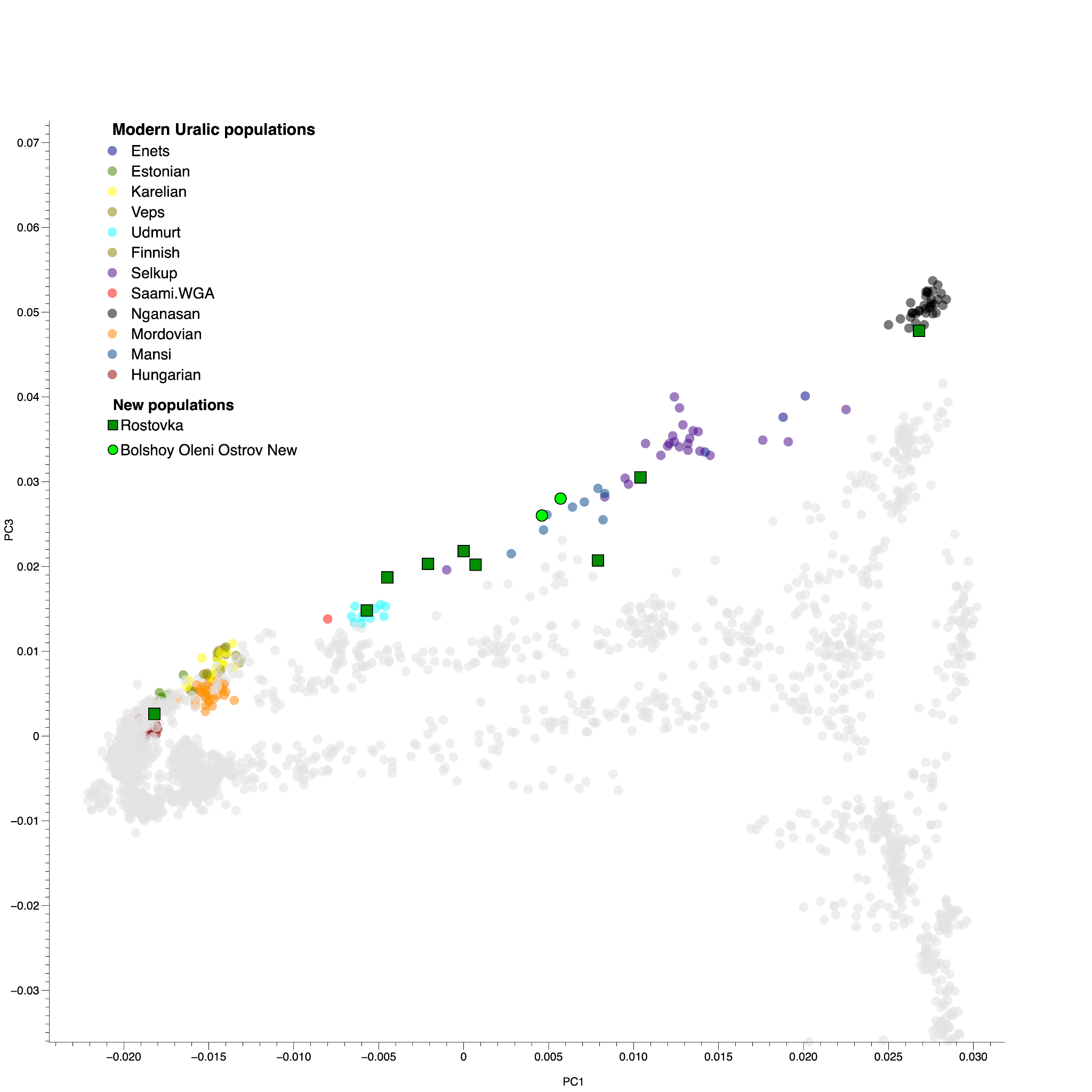

Supplementary Figure 4. PCA (PC1 vs PC3) of present-day Eurasian populations. Modern-day Uralic speakers are highlighted with different colors.

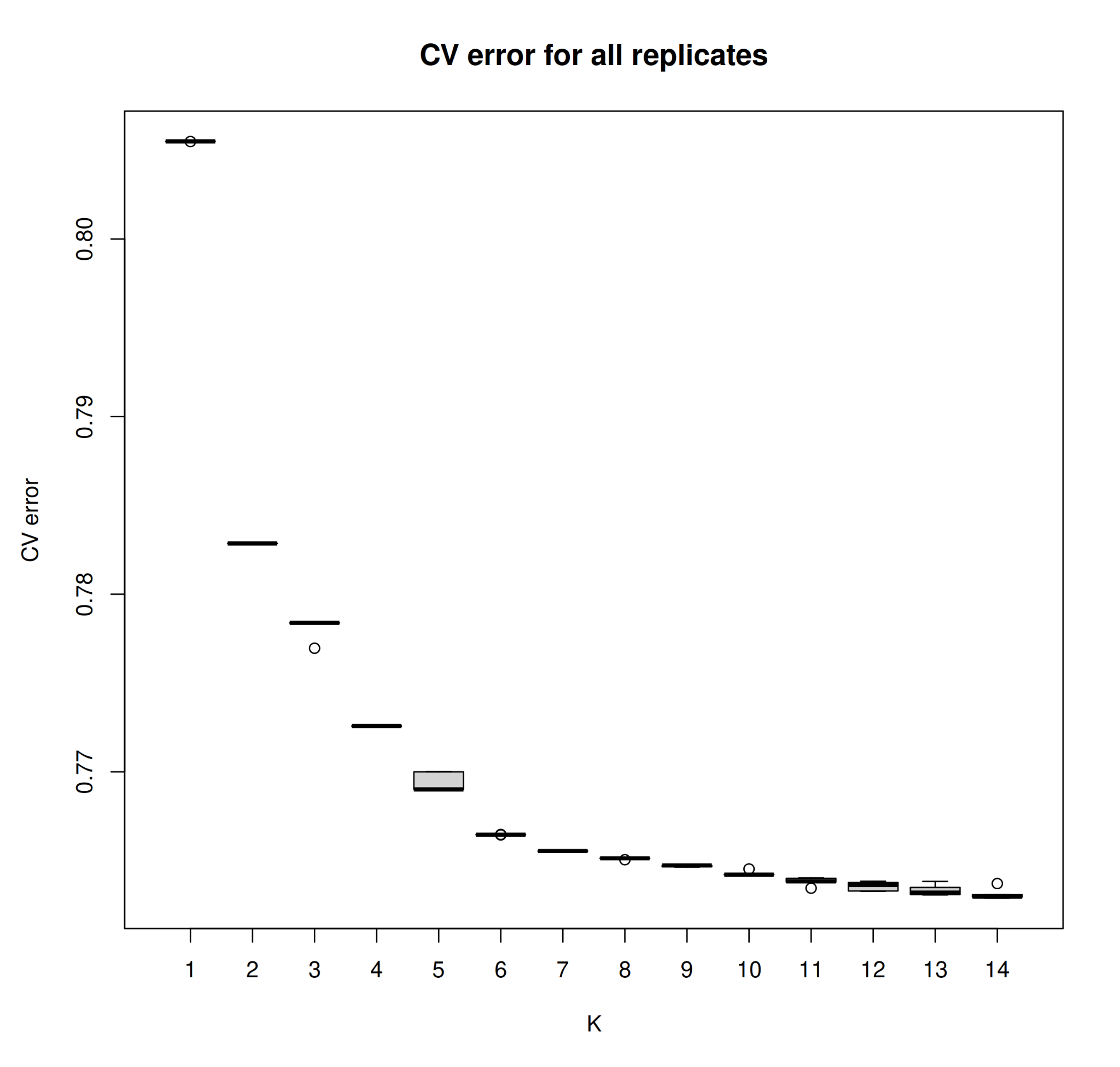

Supplementary Figure 5. Cross-validation (CV) errors for each number of clusters (K) from the unsupervised ADMIXTURE analysis.

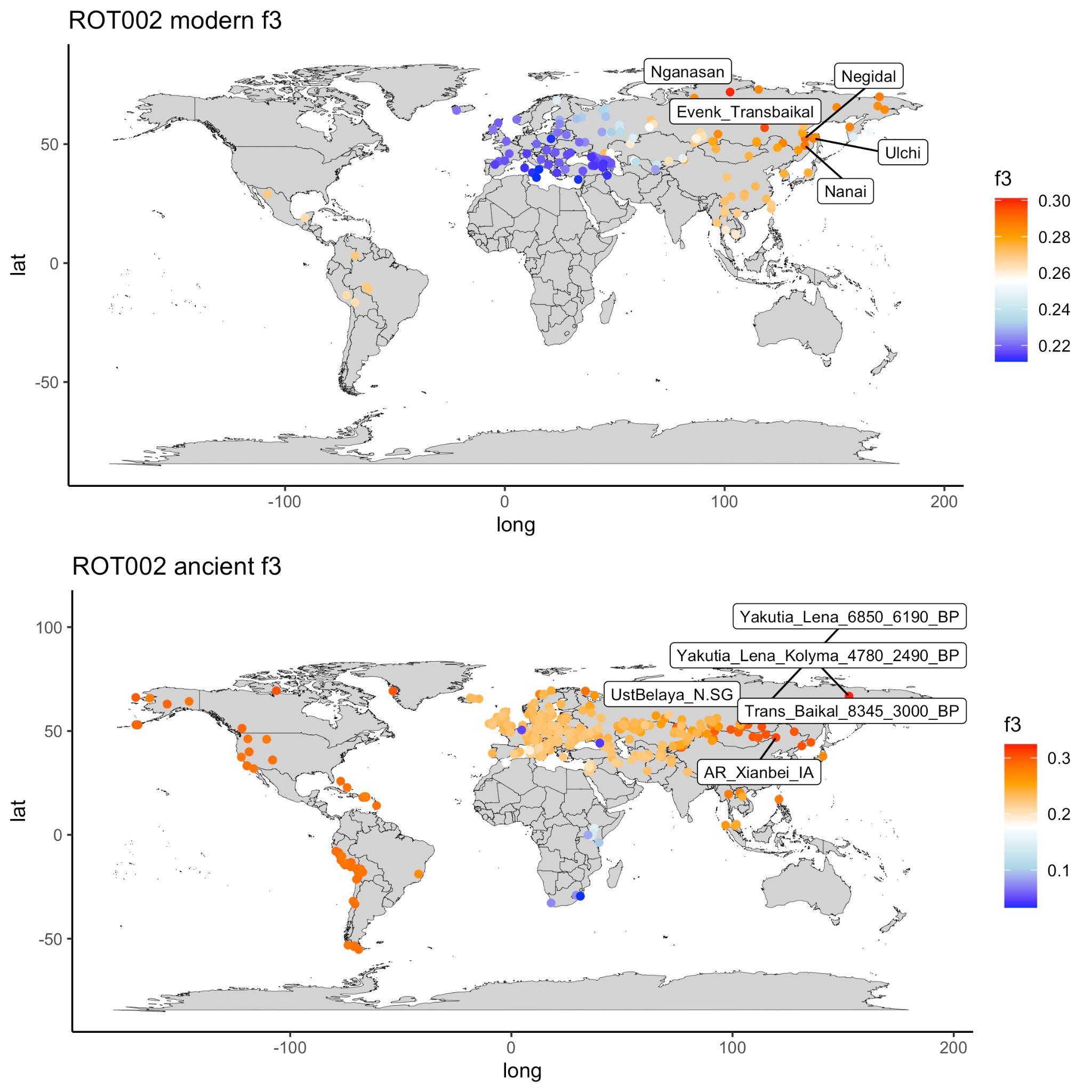
 Supplementary Figure 6A. ROT002 outgroup *f*_3_-statistics with published modern and ancient populations.

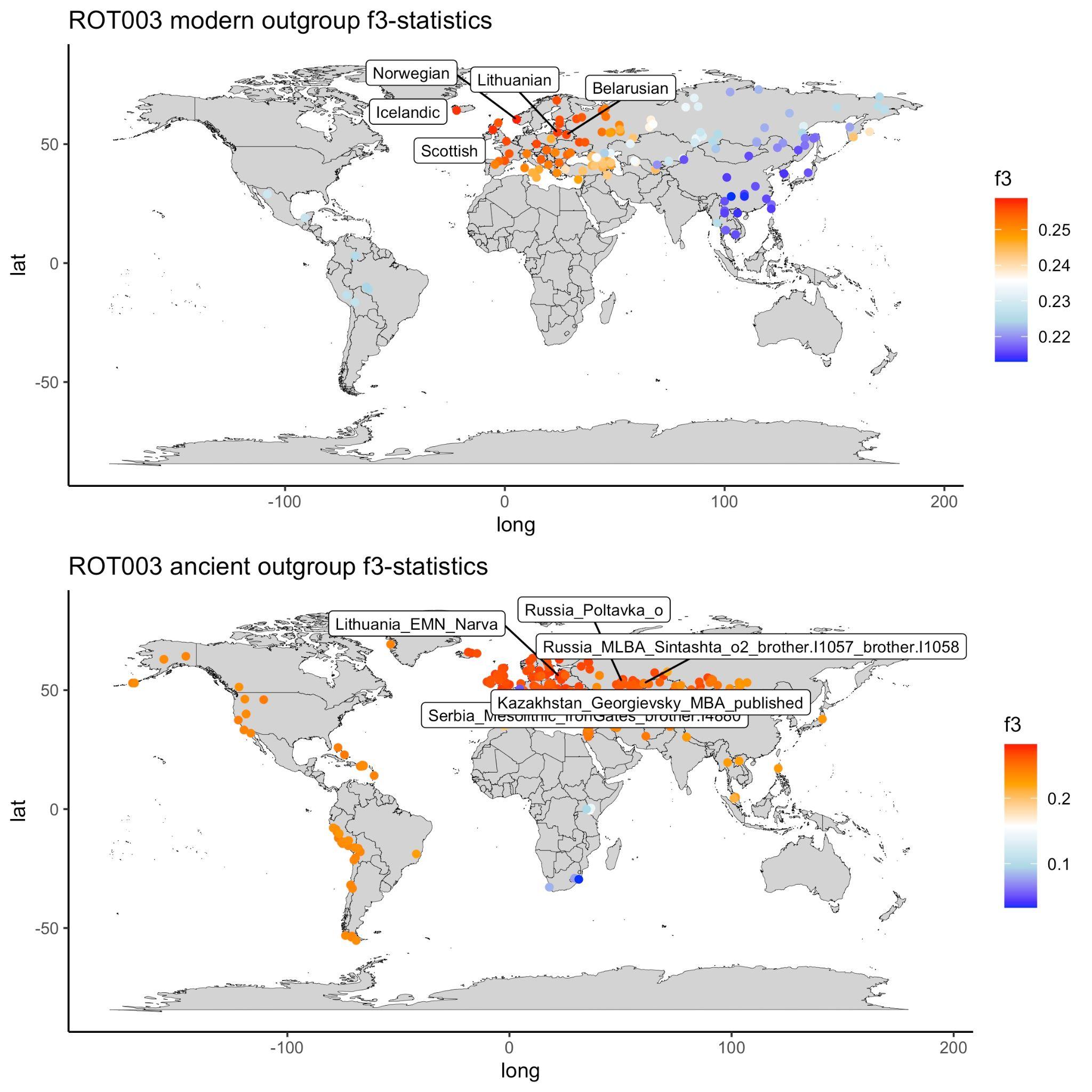
 Supplementary Figure 6B. ROT003 outgroup *f*_3_-statistics with published modern and ancient populations.

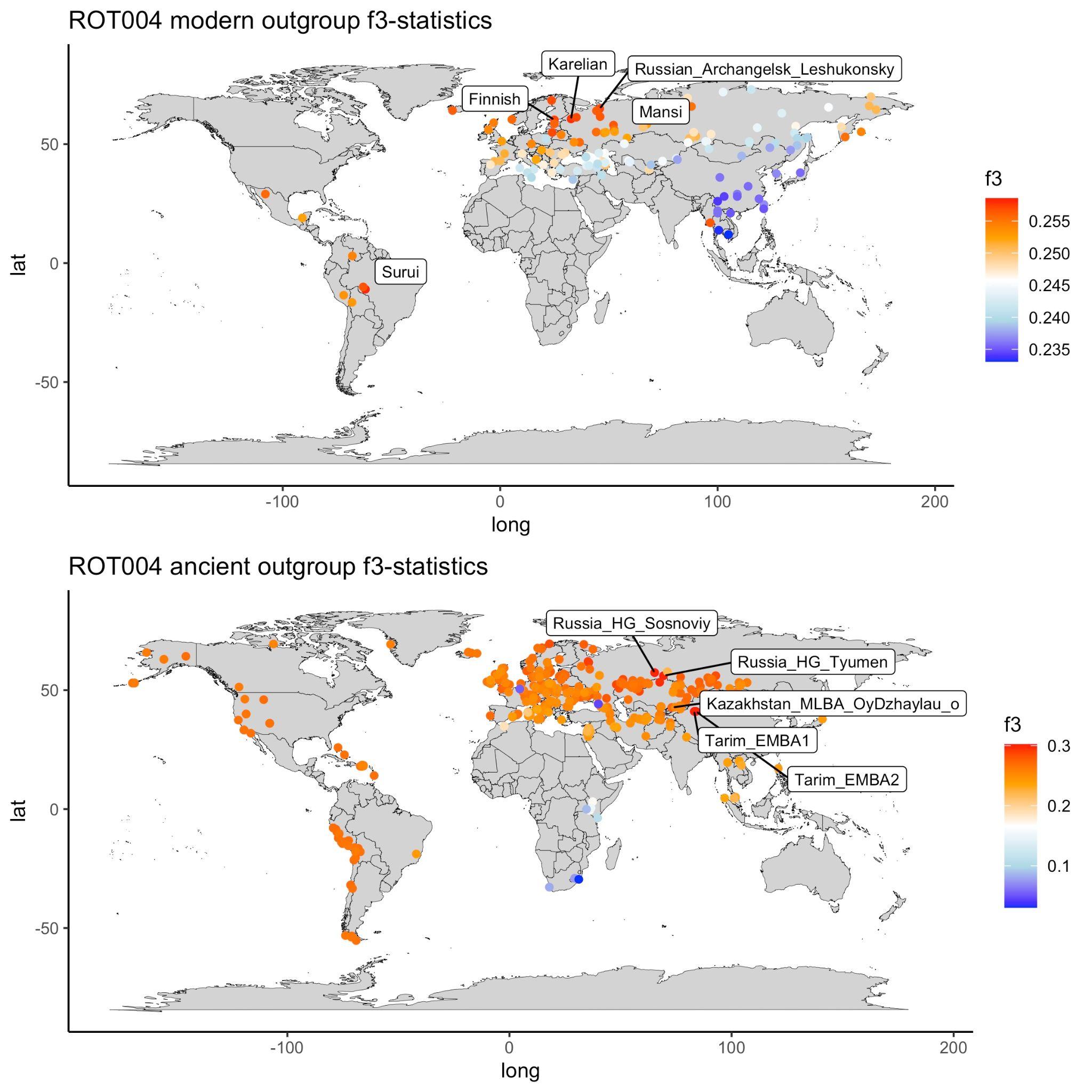
 Supplementary Figure 6C. ROT004 outgroup *f*_3_-statistics with published modern and ancient populations.

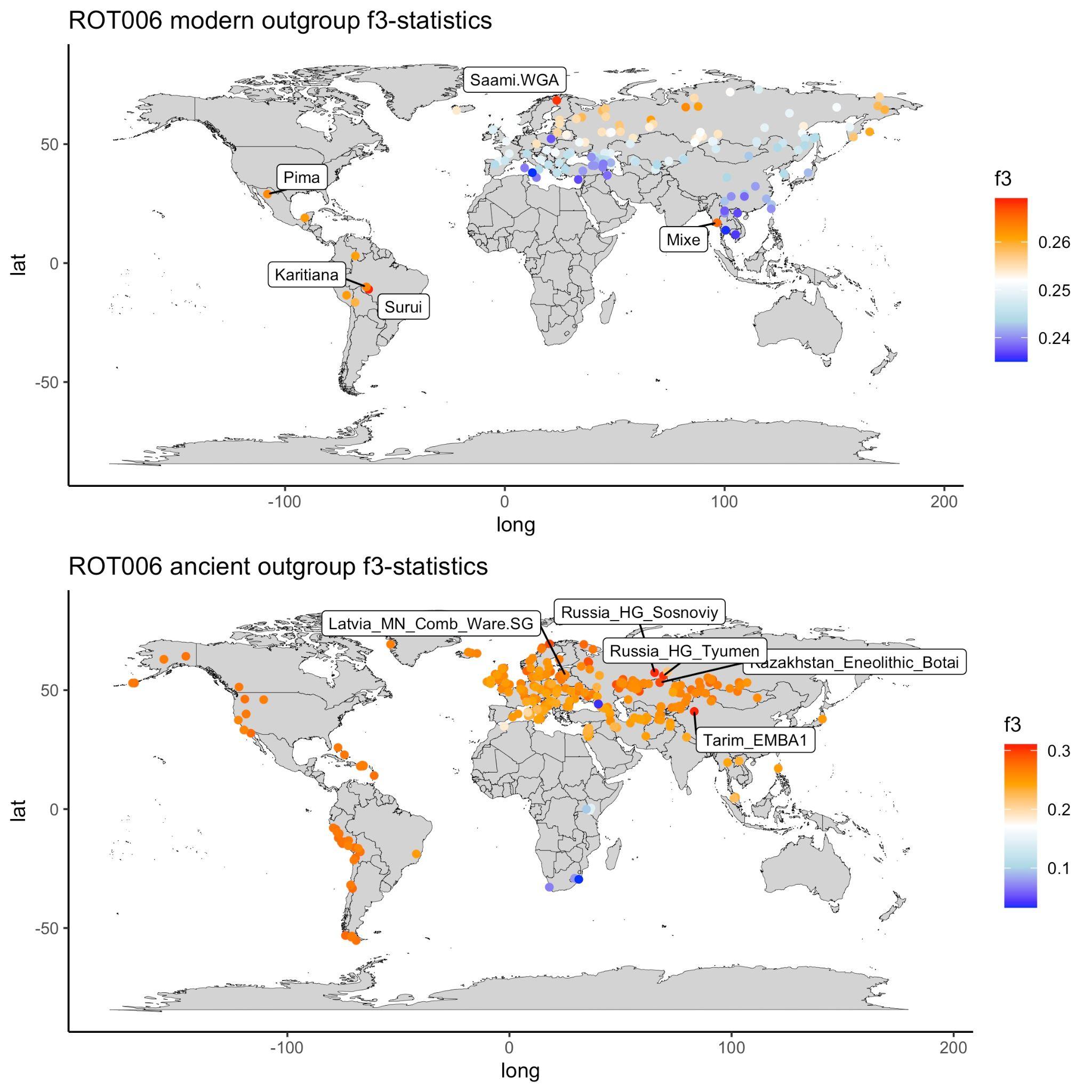
 Supplementary Figure 6D. ROT006 outgroup *f*_3_-statistics with published modern and ancient populations.

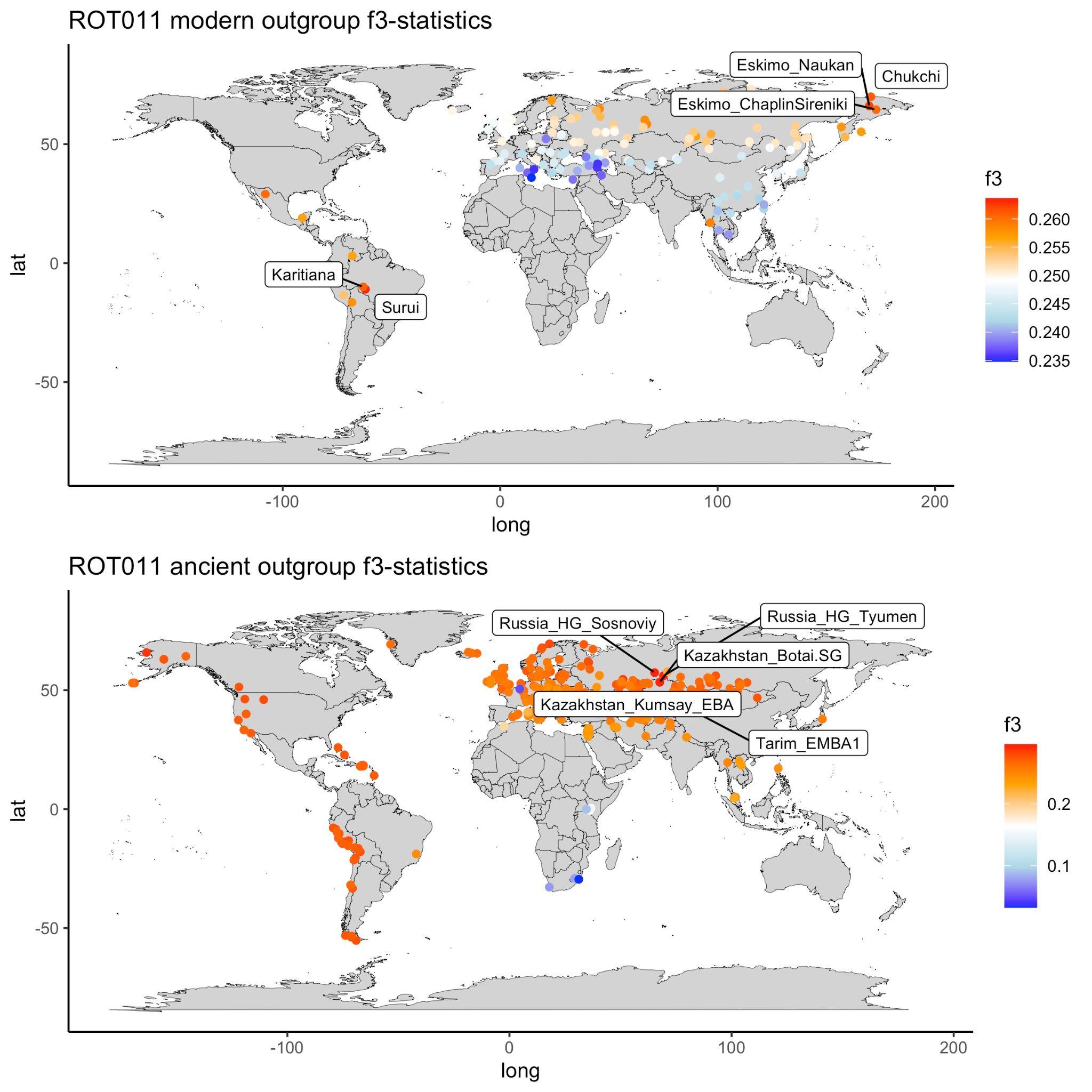
 Supplementary Figure 6E. ROT011 outgroup *f*_3_-statistics with published modern and ancient populations.

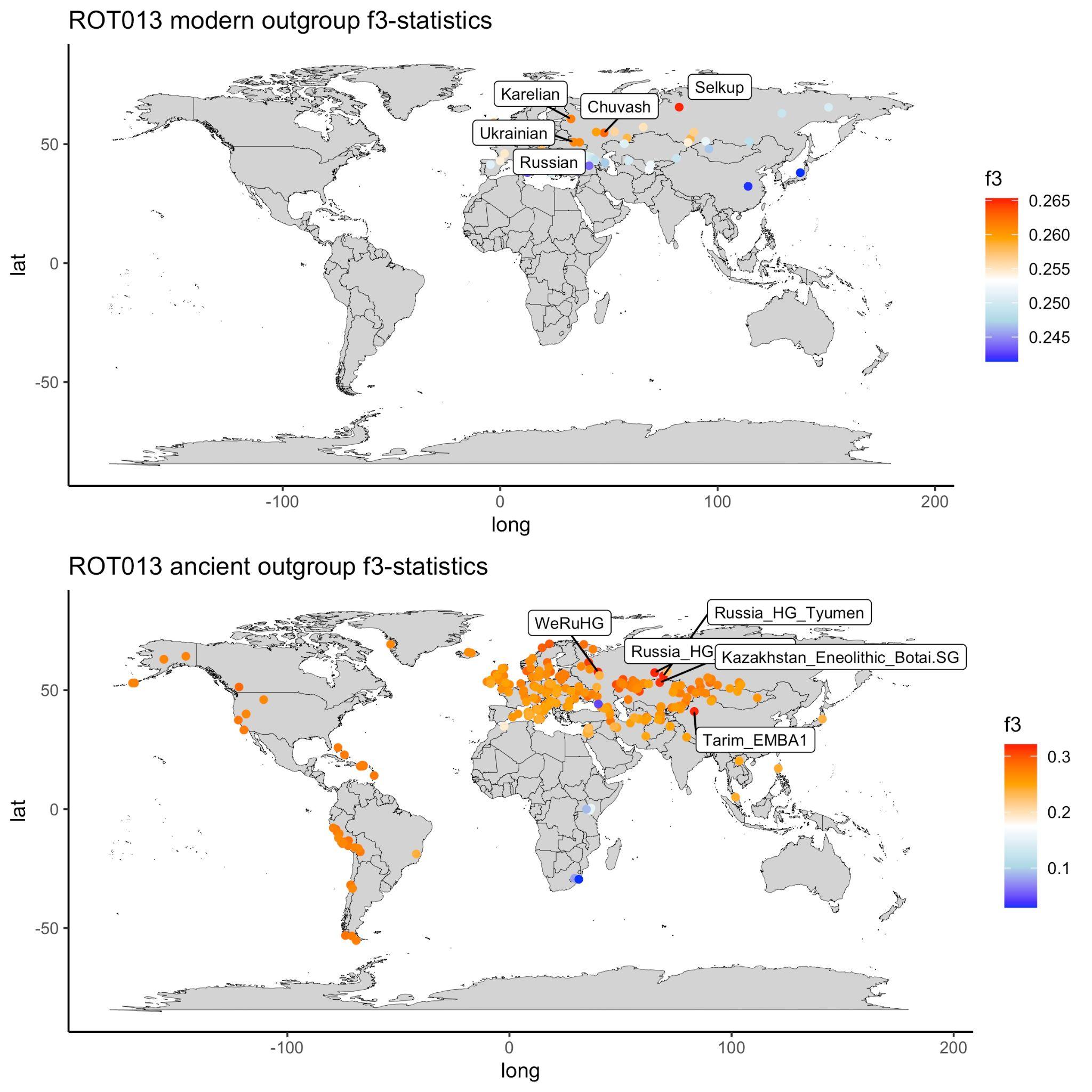
 Supplementary Figure 6F. ROT013 outgroup *f*_3_-statistics with published modern and ancient populations.

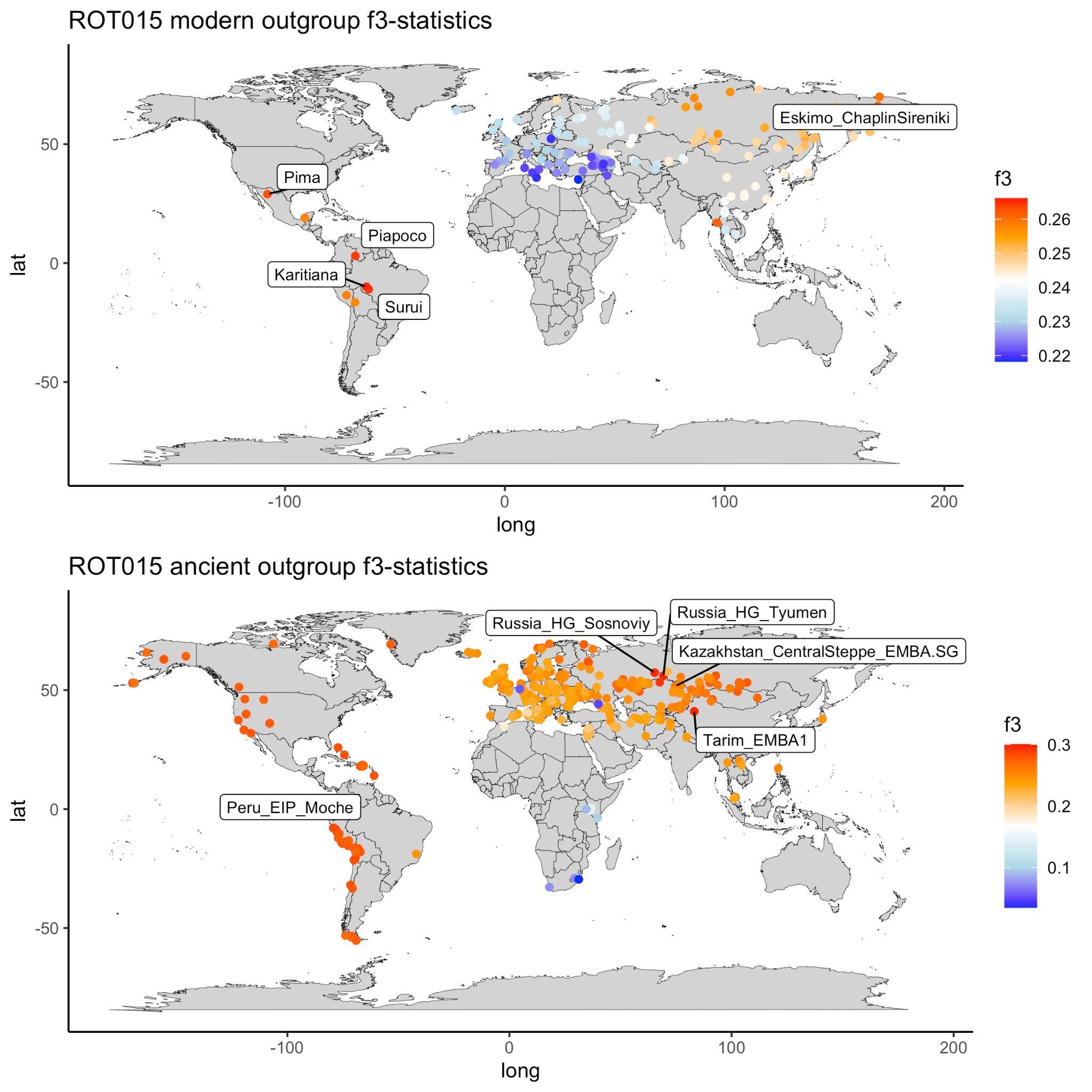
 Supplementary Figure 6G. ROT015 outgroup *f*_3_-statistics with published modern and ancient populations.

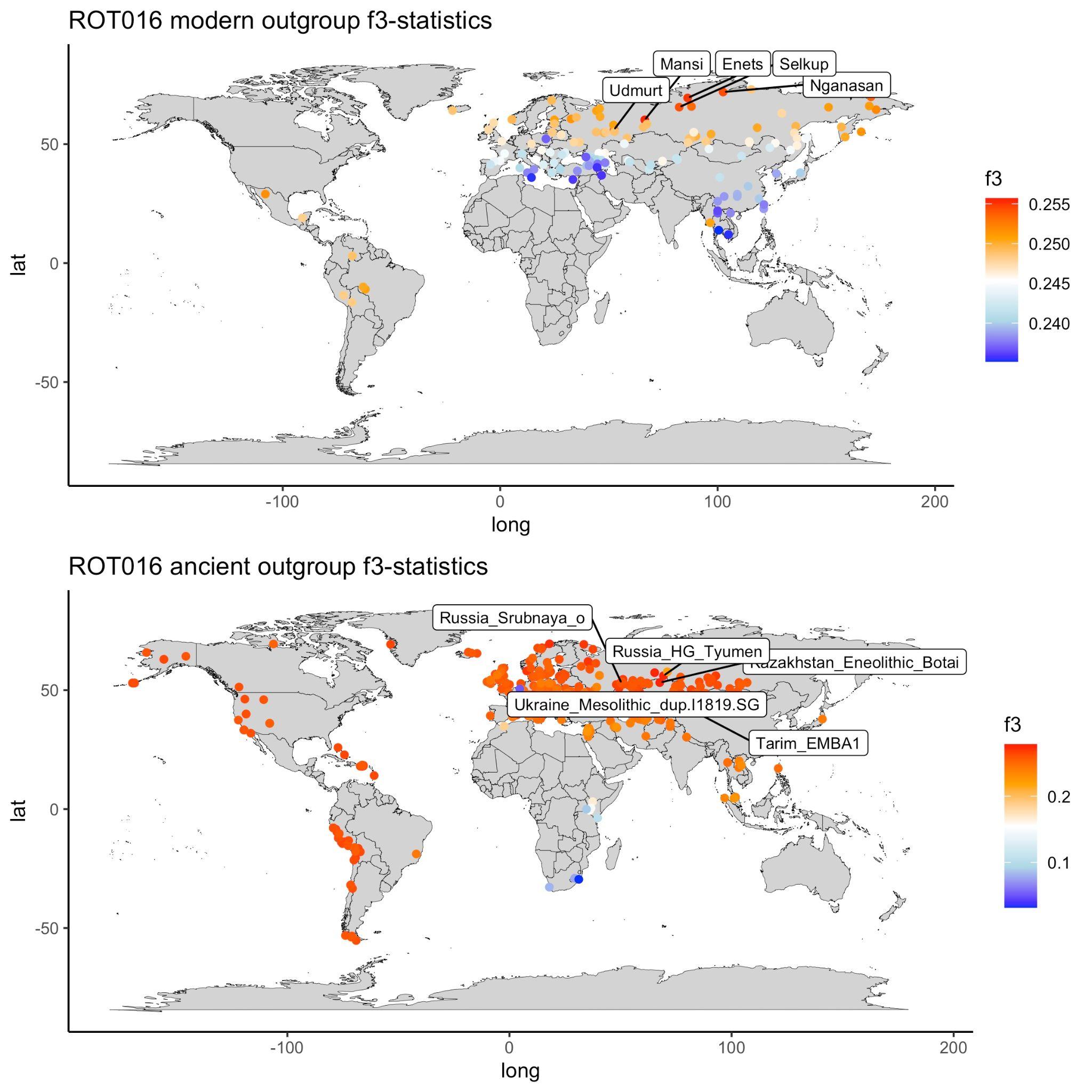
 Supplementary Figure 6H. ROT016 outgroup *f*_3_-statistics with published modern and ancient populations.

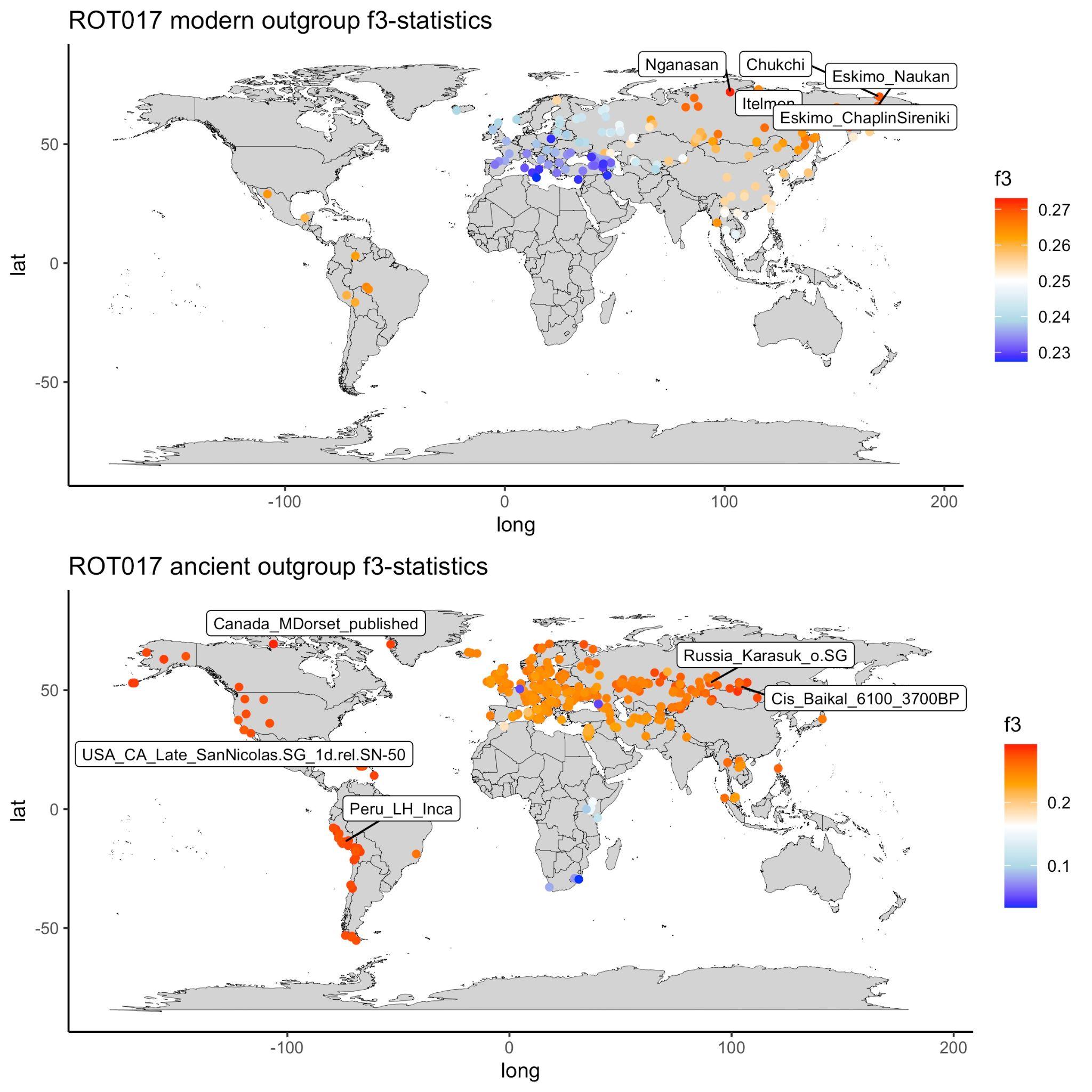
Supplementary Figure 6I. ROT017 outgroup *f*_3_-statistics with published modern and ancient populations.

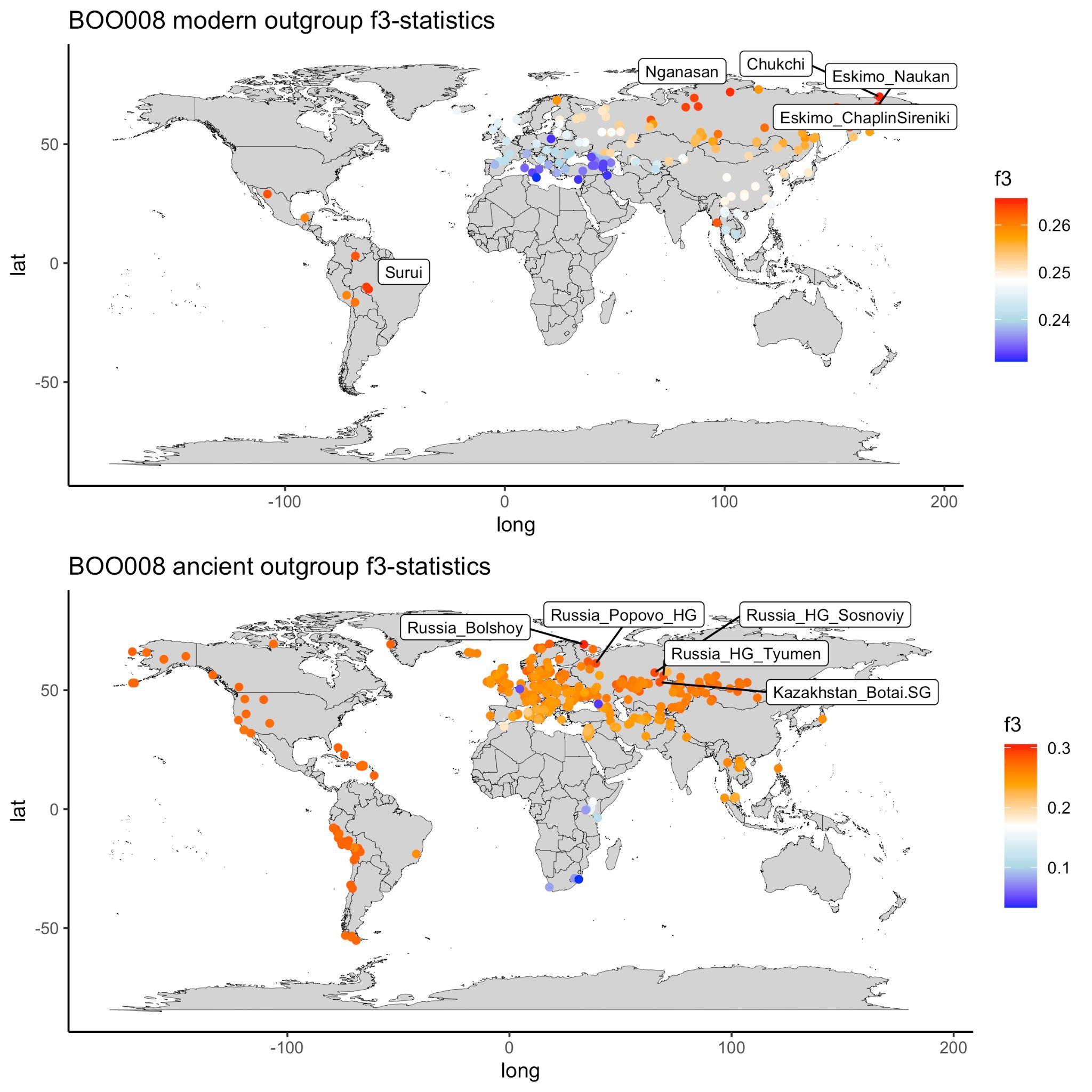
 Supplementary Figure 6J. BOO008 outgroup *f*_3_-statistics with published modern and ancient populations.

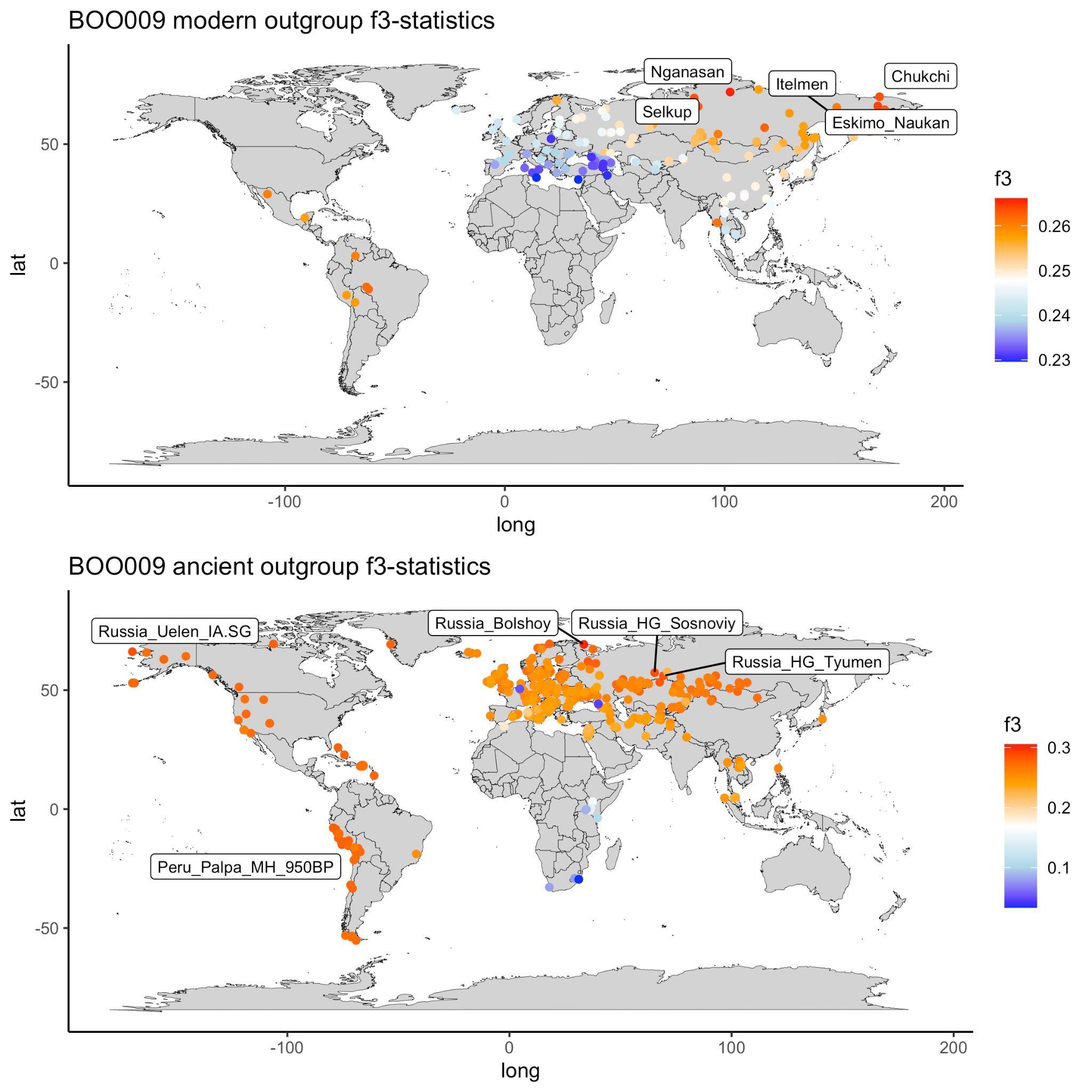
 Supplementary Figure 6K. BOO009 outgroup *f*_3_-statistics with published modern and ancient populations.

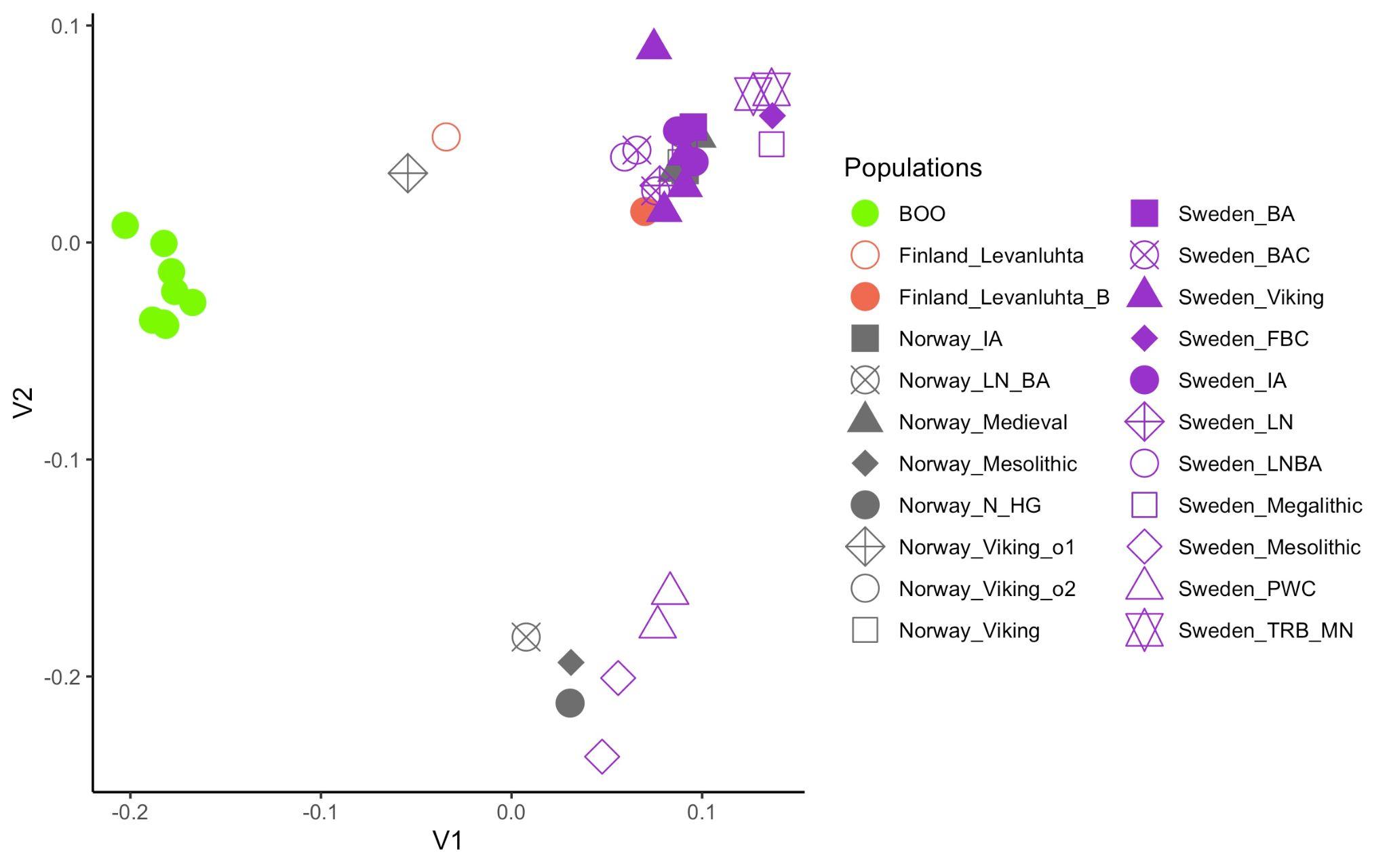

Supplementary Figure 7. MDS plot of pairwise outgroup-*f*_3_ distances. Populations are indicated by shape and color. BOO individuals were analyzed separately. The rest of the f3 models were grouped by population label.

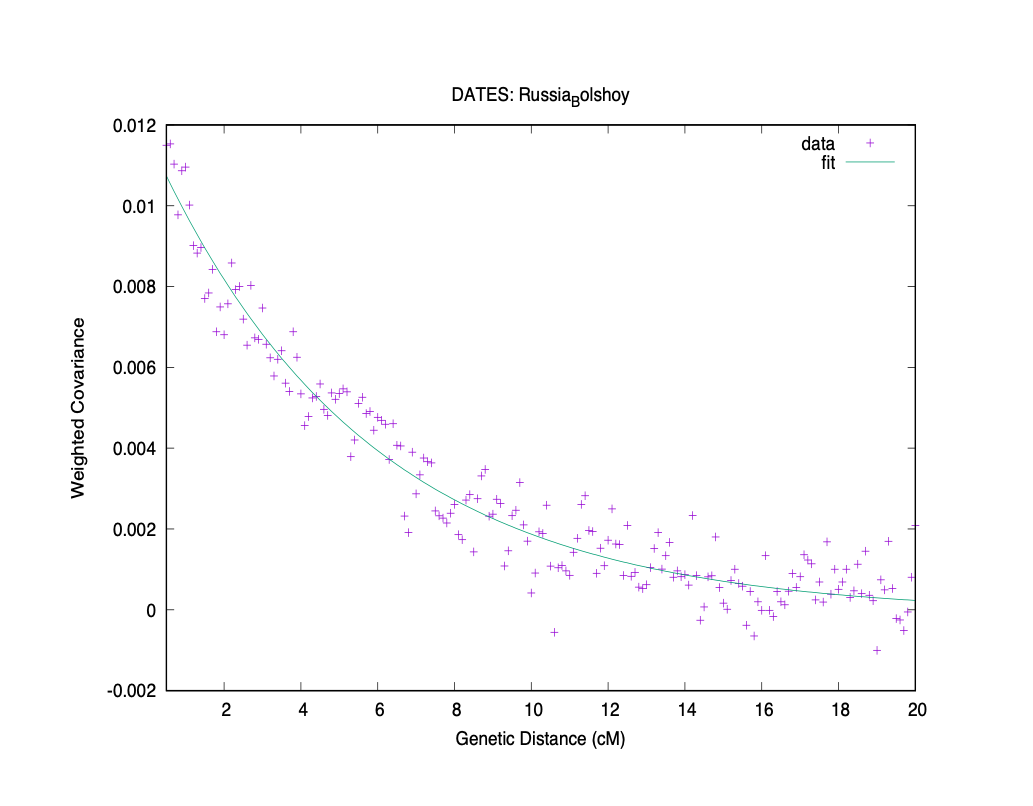
 Supplementary Figure 8. Spatial covariance plot for the DATES admixture dating analysis for BOO when grouped together. Admixture sources are: EEHG and Eastern Siberia LNBA.

**References:**

1.	Chernykh, E. N. & Kuzminykh, S. V. Древняя металлургия Северной Евразии (сейминско-турбинский феномен). https://elibrary.ru/item.asp?id=21143678 (1989).

2.	Matyushenko, V. I. & Sinitsina, G. V. Могильник у д. Ростовка вблизи Омска [Burial Ground near the Village of Rostovka near Omsk]. https://elibrary.ru/item.asp?id=24232508 (1988).

3.	Degtyareva, A. D., and S. V. Kuzminykh. 2011. “Результаты аналитического исследования металлических изделий могильника Сатыга XVI [Results of Analytical Study of Metal Products of Satyga Burial Site XVI].” In *Сатыга XVI: Cейминско-турбинский могильник в таежной зоне Западной Сибири*, 37–44.

4.	Kuzminykh, S. V., V. Y. Lunkov, and L. B. Orlovskaya. 2017. “Результаты рентгенофлуоресцентного анализа: серия 2013-2016 гг [Results of X-Ray Fluorescence Analysis: Series 2013-2016].” In *Аналитические исследования лаборатории естественнонаучных методов.*, 34–60. elibrary.ru.

5.	Lunkov, V. Y., S. V. Kuzminykh, and L. B. Orlovskaya. 2011. “Рентгено-флуоресцентный анализ меди и бронз: серия 2009--2010 гг [X-Ray Fluorescence Analysis of Copper and Bronzes: Series 2009-2010].” In *Аналитические исследования лаборатории естественнонаучных методов*, 116–36. elibrary.ru.

6.	Lunkov, V. Y., S. V. Kuzminykh, and L. B. Orlovskaya. 2013. “Результаты ренгтгено-флуоресцентного анализа: Серия 2011-2013 гг [Results of X-Ray Fluorescence Analysis: Series 2011-2013].” *Аналитические исследования лаборатории естественнонаучных методов*, 56–88.

7.	Lunkov, V. Y., L. B. Orlovskaya, and S. V. Kuzminykh. 2009. “Рентгено-флуоресцентный анализ: начало исследований химического состава древнего металла [X-Ray Fluorescence Analysis: The Beginning of Research into the Chemical Composition of Ancient Metal].” In *Аналитические исследования лаборатории естественнонаучных методов*, 84–110. elibrary.ru.
